# Evaluating the ecological impacts of pesticide seed treatments on arthropod communities in a grain crop rotation

**DOI:** 10.1101/689463

**Authors:** Aditi Dubey, Margaret T. Lewis, Galen P. Dively, Kelly A. Hamby

## Abstract

1. While many studies have investigated non-target impacts of neonicotinoid seed treatments (NSTs), they usually take place within a single crop and focus on specific pest or beneficial arthropod taxa.
2. We compared the impacts of three seed treatments to an untreated control: imidacloprid + fungicide products, thiamethoxam + fungicide products, and fungicide products alone in a three-year crop rotation of full-season soybean, winter wheat, double-cropped soybean and maize. Specifically, we quantified neonicotinoid residues in the soil and in weedy winter annual flower buds and examined treatment impacts on soil and foliar arthropod communities, and on plant growth and yield.
3. Trace amounts of insecticide were found in winter annual flowers of one species in one site year, which did not correspond with our treatments. Although low levels of insecticide residues were present in the soil, residues were not persistent. Residues were highest in the final year of the study, suggesting some accumulation.
4. We observed variable impacts of NSTs on the arthropod community; principle response curve analysis, diversity and evenness values exhibited occasional community disturbances, and treatments impacted the abundance of various taxa. Overall, imidacloprid had a greater effect than thiamethoxam, with the fungicide only treatment also occasionally impacting communities and individual taxa.
5. Pest pressure was low throughout the study, and although pest numbers were reduced by the insecticides no corresponding increases in yield were observed. However, the fungicide products contributed to higher yields in wheat.
6. *Synthesis and applications*. Pesticide seed treatments can disturb arthropod communities, even when environmental persistence and active ingredient concentrations are low. The foliar community in wheat and maize exhibited a trend of increasing disturbance throughout the sampling period, suggesting that recovery from the impacts of NSTs is not always rapid. Our study is among the first to demonstrate that seed applied fungicides alone can disrupt arthropod communities in agroecosystems and highlights the need for further investigation into the impacts of seed applied fungicides.

## 1. Introduction

Declines in arthropod biomass have been documented at multiple locations and are likely linked to habitat loss, climate change, and agrochemical pollutants (Hallmann et al., 2014; Lister & Garcia, 2018). Since their introduction in the 1990s, neonicotinoid insecticides have become the most used insecticide class worldwide, gaining popularity due to their low vertebrate toxicity, systemic nature and versatility of application methods (Nauen, Jeschke, & Copping, 2008). Neonicotinoids are especially popular as seed treatments (NSTs); by 2011, NSTs were used in 79-100% of maize *Zea mays* L. and 34-44% of soybean *Glycine max* L. Merr. planted in the USA (Douglas & Tooker, 2015). When neonicotinoids are applied as NSTs, less than 20% of the active ingredients are taken up by the plant (Alford & Krupke, 2017; Sur & Stork, 2003), instead largely remaining in the soil, where their environmental fate is not fully understood. The half-lives of neonicotinoids in soil vary considerably and they may persist and accumulate for multiple years post planting (Bonmatin et al., 2015). Due to their water solubility, neonicotinoids can also leach into groundwater and run-off into waterbodies; neonicotinoid residues are frequently detected at levels above ecological thresholds in waterbodies that are adjacent to or receive runoff from crop lands (Morrissey et al., 2015). In addition, neonicotinoids may also contaminate non-crop plants. Several studies have found neonicotinoid residues in plants growing near treated fields, but it is difficult to determine whether the active ingredients were taken up from the soil or deposited aerially (Botías et al., 2015; Pecenka & Lundgren, 2015; Stewart et al., 2014). Due to the widespread use, environmental persistence, and mobility of the active ingredients from NSTs, they are common pesticide pollutants.

NSTs pollution can negatively impact many non-target organisms. Although NSTs require relatively low active ingredient concentrations and can reduce non-target exposure due to pesticide drift, they have similar impacts on non-target arthropod abundance as soil and foliar pyrethroid applications (Douglas & Tooker, 2016). Beneficial natural enemies may be exposed to NST active ingredients indirectly by consuming herbivores or directly, either through physical contact or by feeding on plant material or nectar (Gontijo, Moscardini, Michaud, & Carvalho, 2015; Khani, Ahmadi, & Ghadamyari, 2012; Moscardini, Gontijo, Michaud, & Carvalho, 2014; Moser & Obrycki, 2009; Papachristos & Milonas, 2008; Seagraves & Lundgren, 2012). For example, the presence of neonicotinoids in the soil can suppress predatory ground beetles (Coleoptera: Carabidae) through direct contact with active ingredients (Pisa et al., 2015; Simon-Delso et al., 2015), or by ingestion of contaminated prey (Douglas, Rohr, & Tooker, 2015). Work characterizing the impact of neonicotinoids typically focuses on specific pest or beneficial taxa; however, the interconnected arthropod community should also be evaluated as whole. Increased taxon diversity and evenness is associated with reduced pest pressure (Lundgren & Fausti, 2015); therefore, community-level impacts of NSTs could disrupt natural pest control. In maize, clothianidin treated seed disturbed the overall arthropod community after planting, with several beneficial predators decreasing in abundance (Disque, Hamby, Dubey, Taylor, & Dively, 2018). Neonicotinoids can also negatively impact pollinators (Godfray et al., 2014). Because pollinators often rely on non-crop floral resources, uptake by non-crop plants may be an important route of exposure (Botias, David, Hill, & Goulson, 2016). Given the risks associated with NST pollution, the use of NSTs in multiple crops, their potential long-term environmental persistence, and their effects on arthropod communities must all be taken into consideration when evaluating non-target impacts.

In addition to the many risks associated with NSTs, they often provide limited benefits. Active ingredients from NSTs generally remain bioactive in plant tissue for 3-4 weeks post planting, so they only provide protection against early season soil and seedling pests (Alford & Krupke, 2017; Myers & Hill, 2014). Because pest pressure is often not monitored prior to planting, NSTs are frequently used prophylactically, and growers may not recoup the cost of treatment unless significant early season pest pressure occurs (Cox, Cherney, & Shields, 2007; Myers & Hill, 2014; Wilde et al., 2007). The economic benefits of NSTs vary greatly based on region and cropping system and must be evaluated on a case by case basis.

In this study, we evaluated the impacts of repeated use of two popular NSTs [Gaucho 600 (imidacloprid), and Cruiser 5FS (thiamethoxam)] during a three-year grain crop rotation common to the mid-Atlantic United States: full-season soybean, winter wheat, double-cropped soybean and maize. Because commercial NSTs always include fungicides in addition to insecticides, we included a fungicide only treatment as well as an untreated control in order to isolate the impacts of the fungicides from those of the insecticides. To the best of our knowledge, this is among the first studies to quantify the impacts of seed applied fungicides on the arthropod community. The location and concentration of pesticide active ingredients drive non-target effects; therefore, we quantified the persistence of neonicotinoids in the soil and determined whether weedy winter annual flowers uptake residues. We hypothesized that higher levels of neonicotinoid residues would be present in the soil later in the study due to accumulation from multiple crops. Our second object was to evaluate the impacts of pesticide seed treatments on the overall arthropod community and on key arthropod taxa. We anticipated the strongest impacts on the soil community, given the potential for persistence of active ingredients in the soil, and the short activity period in plant tissue. We expected community disturbance early on with recovery during each cropping cycle as observed previously in maize (Disque et al., 2018), but hypothesized that disturbance in the soil community would increase over the course of the study due to potential cumulative impacts of repeated NST use. We also hypothesized that the fungicide only treatment could also impact the arthropod community, due to direct toxicity of seed-applied fungicides towards arthropods (MDA, 2012) or indirect alteration of crop fungal communities. Our final objective was to measure the economic value of the treatments in terms of plant growth metrics and yield, to determine whether the environmental risks of NSTs are justified by economic benefits in mid-Atlantic grain production. We did not expect the insecticide treatment to significantly improve yield because Maryland tends to have low pressure from pests targeted by NSTs; however; neonicotinoids can stimulate plant growth in the absence of pest pressure (Simon-Delso et al., 2015), which could improve growth parameters and yield.

## 2. Materials & Methods

The study was conducted at the Wye Research and Education Center in Queenstown, MD, USA (38°54’02.80” N 76°08’22.06” W) and the Central Maryland Research and Education Center in Beltsville, MD, USA (39°01’08.11” N 76°49’25.10” W) and compared treatments over a three year rotation of four crops at each site. The four treatments were untreated seeds (control), fungicide products alone, fungicide products + imidacloprid insecticide (Gaucho 600; Bayer Crop Science), and fungicide products + thiamethoxam insecticide (Cruiser^®^ 5FS; Syngenta). Full-season soybean was planted in spring 2015, winter wheat in fall 2015, double-cropped soybean in summer 2016, and maize in spring 2017. At each site, four replicate plots of each treatment measuring 9.1m × 15.2m were arranged in a Latin square (Fig. S1). The plot rows were separated by rows of untreated grain that provided space for the planter to turn. Plot columns were separated by three-foot bare strips to delimit plots and facilitate sampling. To determine cumulative effects of repeated treatments, each treatment replicate was planted in the same location for each crop in the rotation. Standard agronomic practices were followed throughout; no foliar fungicides or insecticides were applied. The field at Beltsville was previously planted with untreated soybean, and at Queenstown with neonicotinoid seed treated maize. The seeding rate, variety, and active ingredient rate for each treatment and crop are listed in Tables S1-S2. Due to differences in seeding and application rates, the amount of active ingredient per acre varied slightly between soybean and maize, with wheat concentrations almost double that of the other crops.

### 2.1 Residue analysis

In spring 2016 and 2017, we collected flower buds from winter annual plants growing within the experimental plots for neonicotinoid residue analysis. Winter annual species were chosen based on abundance and attractiveness to pollinators. In 2016, common henbit *Lamium amplexicaule* L. was collected at Beltsville and common chickweed *Stellaria media* L. Vill. at Queenstown. In 2017, we collected common chickweed at Queenstown and both species at Beltsville. Soil was collected for residue analysis before and shortly after soybean and maize were planted in 2015 and 2017, and in March 2016, while wheat was dormant.

Residue samples were sent to the USDA National Science Laboratory (Gastonia, NC, USA) for analysis in 2019, where they were tested for imidacloprid, thiamethoxam, and clothianidin, another popular neonicotinoid that is also a breakdown product of thiamethoxam (Simon-Delso et al. 2015). Further details about material collection and residue analysis are included in section 1.1 of the supporting information.

### 2.2 Arthropod sampling

Throughout the study, the epigeal and soil invertebrate community was measured using sticky cards (3 subsamples per plot), pitfall traps (3 subsamples per plot) and surface litter extractions (4 subsamples pooled into two Berlese funnel extractions per plot). Samples were collected three times during each growing season, and pitfall trap and litter samples were also collected before planting in 2017 maize. In soybean, arthropod abundance in the plant canopy was measured by sweep netting, where 15 sweeps were taken in a straight line through the center of each plot once per season. Samples from one 2015 sweep net imidacloprid replicate at Beltsville and one 2016 sticky card double-cropped soybean sampling date at Queenstown were misplaced prior to processing. We also conducted visual inspections of plants to quantify pest and beneficial arthropods in all crops. The sampling timeline (Tables S3-S6) and further details can be found in section 1.2 of the supporting materials.

### 2.3 Crop sampling

We measured the impact of NSTs on plant growth by recording stand density and plant height in all crops. In wheat, we also counted the number of tillers and measured the Normalized Difference Vegetative Index (NDVI), which can be used to indirectly measure crop biomass (Erdle, Mistele, & Schmidhalter, 2011). We also measured yield at the time of harvest. Details for each crop are included in section 1.3 of the supporting information.

### 2.4 Statistical analysis

For arthropod sampling, taxa were identified to family in most cases, and adults and immatures were combined for all taxa. Insects from the following orders that could not be identified to family were excluded from all analyses: Coleoptera, Diptera, Hemiptera, Hymenoptera and Lepidoptera. After averaging subsamples within each replicate, Shannon Diversity Index (H) and taxa evenness [H/log(n) ratio] were calculated for sticky cards, pitfall traps, litter and sweep nets from each crop using CANOCO 5 (Microcomputer Power, Ithaca, NY, USA).

To characterize the impact of treatment over time, arthropod community composition was analyzed in CANOCO 5 using Principal Response Curve Analysis (PRC) for pitfall traps, litter extraction and sticky card data for each crop (Disque et al. 2018). Briefly, PRC multivariate analysis is based on Redundancy Analysis (Van den Brink & Ter Braak J. F., 1999), with adjustments for the change in community response over time. In our study, total abundances for each taxon were averaged over subsamples within a replicate plot for each site prior to analysis. Taxa where the sum of individuals across sampling dates and sites for a crop was less than one were excluded from the PRC. For each crop and sample type, the date*treatment interaction term was used as an explanatory variable, and date and the site*replicate interaction were used as covariates to restrict data shuffling. Canonical coefficients were generated for each date and plotted over time to evaluate the community response to the treatments relative to the untreated control; the control is plotted along the horizontal axis (representing time), with the magnitude (represented by canonical coefficients plotted on the vertical axis) and shape of curves representing the deviation of treatments from the control. The analysis also generates taxon-specific weights for the individual taxa that exhibit the strongest effects; taxa with high positive weights are more likely to follow the pattern depicted in the PRC, while taxa with high negative weights exhibit an opposite response. A Monte-Carlo permutation procedure with N=499 was used to test the null hypothesis that the canonical coefficients of the treatment response equaled zero for all sampling times, and to calculate a Pseudo-F statistic (Disque et al., 2018). Because sweep net samples were conducted on a single date, captures were analyzed using RDA (Van den Brink & Ter Braak J. F., 1999). Ants (Hymenoptera: Formicidae) were excluded from PRC, diversity and evenness for sticky cards, pitfall traps and litter due to their highly clumped distribution on the ground, but were included in RDA, diversity and evenness for sweep net sampling.

Analysis of variance (Proc Mixed, SAS 9.4, SAS Institute Inc., Cary, NC, USA) was used to evaluate treatment effects for Shannon Index, evenness, stand count, plant height, NDVI, tiller count, yield and abundances of key taxa. For each crop and sample type, treatment was included as an explanatory variable, and site, row and column were random blocking factors, with row and column nested within site. For data collected on multiple dates, date and date*treatment interactions were also included as fixed effects. The date*treatment interaction was dropped when not significant. Before analysis, residual plots and the Shapiro-Wilk’s W test were performed to examine data normality and homogeneity of variances, and appropriate transformations or variance groupings were applied as needed. When P<0.10 for the treatment effect, contrasts were used to compare the fungicide, imidacloprid and thiamethoxam treatments to the control.

## 3. Results

### 3.1 Residue analysis

#### 3.1.1 Winter annual flowers

The detection level was 10 ppb for imidacloprid, 5 ppb for thiamethoxam and 30 ppb for clothianidin in flowers. In 2016, neonicotinoid residues were not found in any samples. In 2017, trace amounts (<10ppb) of imidacloprid were found in five of the chickweed samples from Beltsville, specifically two control samples and one from each of the other treatments. Detections did not occur in a spatial pattern.

#### 3.1.2 Soil

In soil the detection level was 5ppb for imidacloprid, 10ppb for thiamethoxam and 15ppb for clothianidin. Before planting in 2015, low levels (≤10ppb) of imidacloprid were present in several replicates at Beltsville and no residues were present at Queenstown (Table S7). Similar levels were detected after treated soybean was planted, and trace amounts of thiamethoxam and clothianidin were found in one thiamethoxam and one imidacloprid treated replicate. Only one thiamethoxam replicate contained detectable residues at Queenstown. In 2016, during wheat dormancy, 7ppb of imidacloprid was found in the imidacloprid treated plots from Beltsville, with trace amounts in the other plots from Beltsville and the imidacloprid plots from Queenstown. Before maize was planted in 2017, low levels of imidacloprid were present in both imidacloprid sample replicates, and one control and thiamethoxam sample replicate at Beltsville. After maize was planted, imidacloprid was detected across multiple treatments at Beltsville, and in the imidacloprid treated plots at Queenstown, with higher levels (≥10ppb) present in the imidacloprid treated plots at both sites. Thiamethoxam was detected in both thiamethoxam replicates (15-16ppb) at Queenstown, and thiamethoxam (17ppb) and clothianidin (23ppb) were found in one thiamethoxam replicate from Beltsville.

### 3.2 Arthropod sampling

The date by treatment interaction was not significant for arthropod and crop analyses. Therefore, the interaction term was removed from the model and we present information solely for the treatment effect.

#### 3.2.1 Effects of seed treatments on the overall community over time

##### 3.2.1.1 Diversity and evenness

###### Pitfall traps

Activity density of arthropods collected using pitfall traps was measured during three one-week periods. After averaging subsamples and removing unidentifiable individuals and ants (Formicidae), we collected 53,359 individuals across sites and crops (Table S8). Shannon diversity indices were not impacted for any crop: 2015 soybean (F_3,77_=0.43, P=0.729), 2015-2016 wheat (F_3,77_=0.95, P=0.420), 2016 double cropped soybean (F_3,77_=0.75, P=0.528), pre-planting 2017 maize (F_3,15_=1.23, P=0.333), post-planting 2017 maize (F_3,77_=0.79, P=0.502). Similarly, evenness was also not impacted: 2015 soybean (F_3,77_=0.30, P=0.828), 2015-2016 wheat (F_3,77_=0.78, P=0.509), 2016 double cropped soybean (F_3,77_=0.29, P=0.831), pre-planting 2017 maize (F_3,15_=0.57, P=0.640), post-planting 2017 maize (F_3,77_=0.60, P=0.617).

###### Litter extraction

Impacts on arthropod abundance were measured by litter extraction three times in each crop. After averaging subsamples, a total of 69,332 individuals were identified from across crops and sites (Table S9). Shannon diversity indices for litter communities were generally similar across treatments in the first two years: 2015 soybean (F_3,77_=0.14, P=0.937), 2015-2016 wheat (F_3,77_=2.83, P=0.044, no contrast differences), and 2016 double cropped soybean (F_3,77_=0.67, P=0.571). However, in 2017 maize both insecticide treatments reduced diversity (F_3,77_=3.90, P=0.031) relative to the control pre-planting (Table 1). Post-planting (F_3,77_=3.97, P=0.011), only imidacloprid reduced diversity. Litter community evenness was also not impacted until the last year of the study: 2015 soybean (F_3,77_=0.25, P=0.863), 2015-2016 wheat (F_3,77_=1.56, P=0.206), and 2016 double cropped soybean (F_3,77_=0.78, P=0.507). In 2017, taxa evenness was reduced by imidacloprid in pre-planting plots (F_3,15_=2.53, P=0.096) and by all pesticide seed treatments in post-planting plots (F_3,77_=3.76, P=0.014) (Table 1).

**Table 1.** Treatment means for Shannon Index and evenness values for litter extraction (LE), sticky card (SC) and sweep net (SN) arthropod communities. Analysis of variance was used with date and treatment as fixed effects and location, row (location) and column (location) as random effects. Contrasts were used to compare the fungicide (FUN), imidacloprid (IMI) and thiamethoxam (THI) treatments to the control (CON), * indicates P<0.05, ** indicates P<0.01, *** indicates P<0.001 and N.S. indicates no significance. FS = full season and DC = double cropped soybean.

| Metric | Crop | Sample Type | Treatment Mean ± S.E. | | | |
| --- | --- | --- | --- | --- | --- | --- |
|  |  |  | CON | FUN | IMI | THI |
| Shannon Index | FS Soybean (2015) | SC | $2.02 \pm 0.06$ | $1.97 \pm 0.07$ <sup>N.S.</sup> | $1.94 \pm 0.07$ * | $2.04 \pm 0.07$ <sup>N.S.</sup> |
| | Winter Wheat (2015-2016) | LE | $1.27 \pm 0.05$ | $1.36 \pm 0.05$ <sup>N.S.</sup> | $1.17 \pm 0.07$ <sup>N.S.</sup> | $1.28 \pm 0.05$ <sup>N.S.</sup> |
| | DS Soybean (2016) | SN | $2.24 \pm 0.11$ | $2.13 \pm 0.05$ <sup>N.S.</sup> | $2.09 \pm 0.07$ <sup>N.S.</sup> | $1.90 \pm 0.08$ ** |
| | Maize Pre-Plant (2017) | LE | $1.71 \pm 0.10$ | $1.63 \pm 0.07$ <sup>N.S.</sup> | $1.3 \pm 0.14$ * | $1.45 \pm 0.06$ * |
| | Maize Post-Plant (2017) | LE | $1.47 \pm 0.09$ | $1.42 \pm 0.11$ <sup>N.S.</sup> | $1.22 \pm 0.07$ ** | $1.36 \pm 0.10$ <sup>N.S.</sup> |
| Evenness | FS Soybean (2015) | SN | $0.67 \pm 0.06$ | $0.77 \pm 0.04$ * | $0.74 \pm 0.05$ * | $0.77 \pm 0.05$ ** |
| | Winter Wheat (2015-2016) | SC | $0.72 \pm 0.03$ | $0.72 \pm 0.02$ <sup>N.S.</sup> | $0.74 \pm 0.02$ <sup>N.S.</sup> | $0.75 \pm 0.03$ * |
| | DC Soybean (2016) | SN | $0.83 \pm 0.04$ | $0.84 \pm 0.02$ <sup>N.S.</sup> | $0.83 \pm 0.03$ <sup>N.S.</sup> | $0.77 \pm 0.04$ <sup>N.S.</sup> |
| | Maize Pre-Plant (2017) | LE | $0.74 \pm 0.04$ | $0.71 \pm 0.03$ <sup>N.S.</sup> | $0.61 \pm 0.07$ * | $0.65 \pm 0.05$ <sup>N.S.</sup> |
| | Maize Post-Plant (2017) | LE | $0.73 \pm 0.03$ | $0.61 \pm 0.04$ * | $0.58 \pm 0.03$ ** | $0.61 \pm 0.04$ * |

###### Sticky cards

Arthropod activity density ∼8cm above the ground was measured during three one-week intervals using sticky cards. Date from the second sampling date at Queenstown for double-cropped soybean could not be included because the cards were misplaced before processing. After averaging subsamples, 34,413 individuals were identified from across crops and sites (Table S10). Shannon diversity indices were not impacted by the pesticide treatments for most crops: 2015-2016 wheat (F_3,77_=1.11, P=0.351), 2016 double cropped soybean (F_3,61_=1.59, P=0.201), 2017 maize (post-planting) (F_3,77_=1.96, P=0.127). However, in 2015 full-season soybean (F_3,77_=2.36, P=0.078), sticky card captures were less diverse in imidacloprid plots than control plots. Treatments also did not impact taxa evenness for most crops: 2015 soybean (F_3,77_=0.73, P=0.537), 2016 double cropped soybean (F_3,77_=1.64, P=0.189), 2017 maize (post-planting) (F_3,77_=0.27, P=0.847); except in wheat (F_3,77_=2.28, P=0.082), where the thiamethoxam treatment exhibited higher taxa evenness (Table 1).

###### Sweep net

Sweep nets were used to collect canopy-dwelling taxa in full-season and double-cropped soybean, with 3,892 individuals identified (Table S11). In full-season soybean, diversity was not impacted by the pesticide treatments (F_3,14_=1.78, P=0.197), while in double-cropped soybean (F_3,15_=3.6, P=0.039), the thiamethoxam treatment exhibited lower arthropod diversity than the control (Table 1). Taxa evenness was lower than the control in all three treatments in 2015 (F_3,14_=3.95, P=0.031), but not 2016 (F_3,15_=3.18, P=0.055, no contrast differences).

##### 3.2.1.2 Community disturbances

###### Pitfall traps

Pitfall trapped communities exhibited no disturbance by the pesticide treatments over time for any crop: 2015 full-season soybean (Pseudo-F=0.1, P=0.924), 2015-2016 winter wheat (Pseudo F=0.2, P=0.712), 2016 double-cropped soybean (Pseudo-F=0.2, P=0.814), 2017 maize (Pseudo F=0.2, P=0.278) (Fig. S2).

###### Litter extraction

Litter communities were similarly not disturbed across sampling dates by the pesticide treatments: 2015 full-season soybean (Pseudo-F=0.2, P=0.946), 2015-2016 winter wheat (Pseudo-F=0.2, P=0.976), 2016 double-cropped soybean (Pseudo-F=0.3, P=0.064), 2017 maize (Pseudo-F=0.5, P=0.198) (Fig. S3). Though the insecticide treatment canonical coefficients were often below the control on the first sampling date, community responses varied.

###### Sticky cards

Although sticky card communities were not impacted by pesticide treatments in 2015 full-season (Pseudo-F=0.2, P=0.356) and 2016 double-cropped (Pseudo-F=0.3, P=0.198) soybean, increasing insecticide treatment community declines were observed over the sampling dates in wheat (Pseudo-F=0.5, P=0.002), with all pesticides causing increasing declines over time in maize (Pseudo-F=0.3, P=0.078) (Fig. 1).

**Fig. 1.**
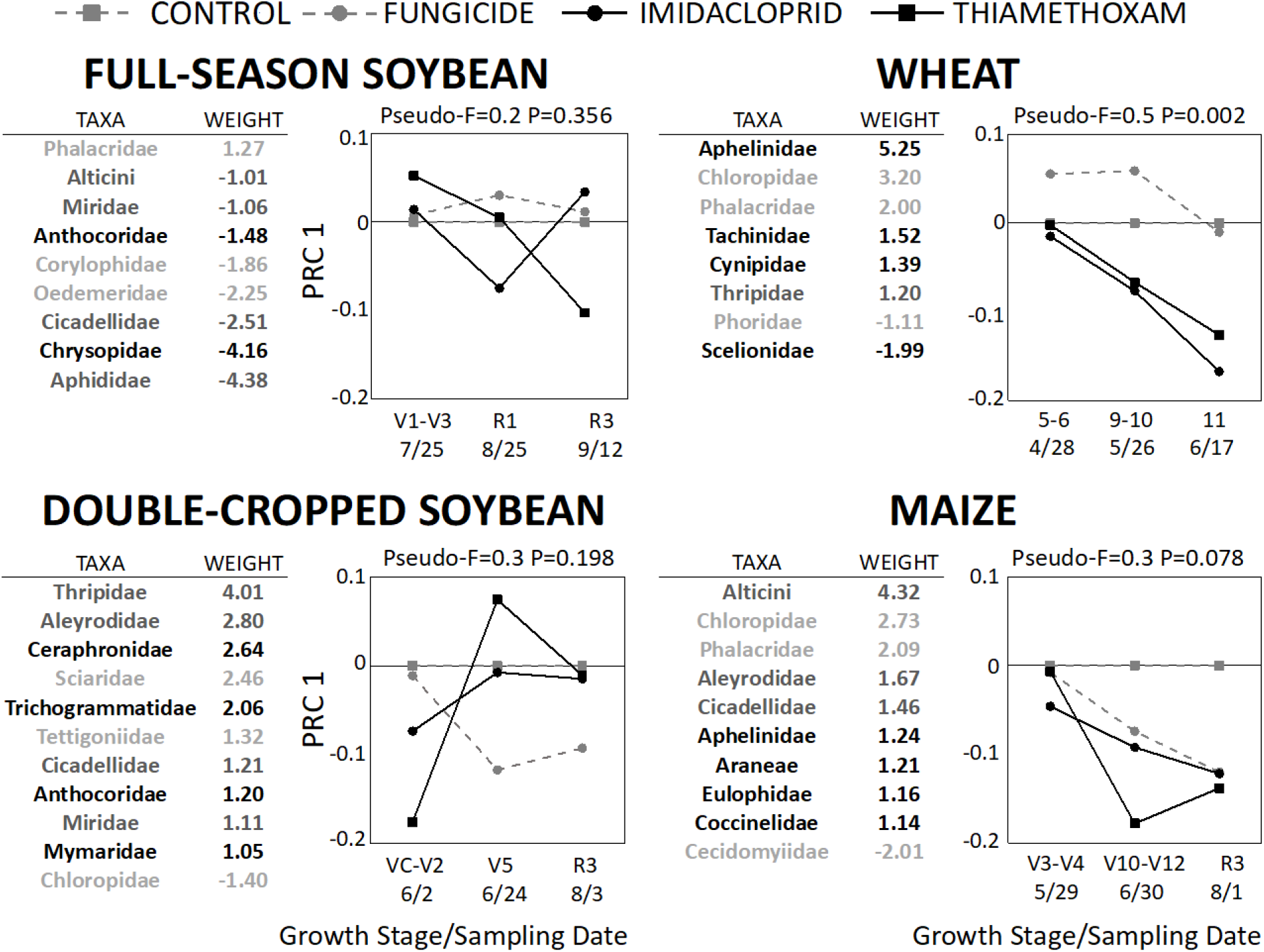
Principal Response Curve analysis of sticky card data for all crops. Date*treatment served as the explanatory variable, with date and site*replicate used as covariates. Subsamples were averaged by taxa for each replicate, and only taxa with overall means greater than one were included. Ants (Formicidae) were also excluded due to their highly clumped distribution. A Monte-Carlo permutation procedure with N=499 was used to calculate the Pseudo-F statistic. Taxon weights indicate which groups most contributed to the observed community response. Higher positive weights indicate that taxon abundances in the treated plots followed the trend depicted by the response curve, whereas higher negative values indicate the opposite. Taxon weights between −1 and 1 were excluded due to weak response or lack of correlation with the trends shown. Beneficial groups are shown in black, economic pests in dark gray, and other groups in light gray.

###### Sweep net

Sweep net collected arthropods were not impacted by pesticide treatments in full-season soybean (First axis Pseudo-F=0.4, P=0.412); however, multiple beneficial taxa (e.g., Coccinellidae, Anthocoridae, Araneae) were reduced by the insecticide treatments in double-cropped soybean (First axis Pseudo-F=0.9 P=0.004) (Fig 2).

**Fig. 2.**
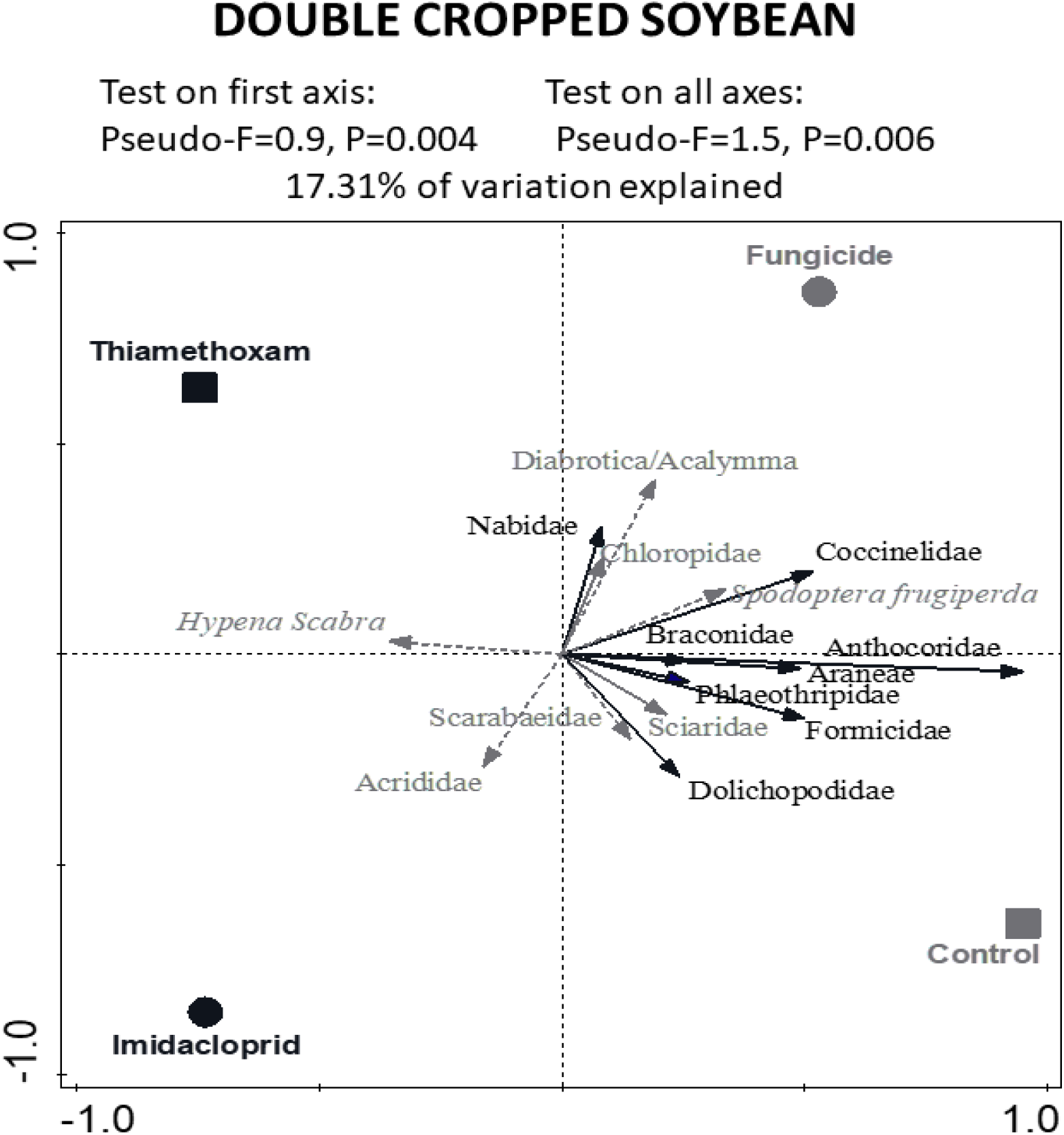
Redundancy Analysis of sweep net data from 2016 double cropped soybean. Treatment served as the explanatory variable and the site*replicate interaction was used as a covariate. The horizontal axis is the first axis. Only the 15 taxa that most contributed are shown. A Monte-Carlo permutation procedure with N=499 was used to calculate a Pseudo-F statistic. Beneficial groups are shown in black, economic pests in gray with dotted lines, and other groups in gray with solid lines.

#### 3.2.2 Effects of seed treatments on individual taxa within crops

##### 2015 Full-season soybean

Visual scouting revealed fewer predatory thrips (Phlaeothripidae, F_3,15_=4.46, P=0.020) within both insecticide treated plots; however, only thiamethoxam reduced leafhoppers (Cicadellidae, F_3,15=_8.77, P=0.001) and pestiferous thrips (Thripidae, F_3,15_=8.21, P=0.002) (Table 2).

**Table 2.** Treatment means of individual taxa from pitfall traps (PT), litter extraction (LE), sticky cards (SC), sweep nets (SN) and visual counts (VC). Data was averaged across treatment dates for sticky card, pitfall trap, litter and soybean and wheat visual count samples; in maize visual counts, each date was analyzed separately due to differences in sampling methodology. Analysis of variance was used with date and treatment as fixed effects and location, row (location) and column (location) as random effects. Contrasts were used to compare the fungicide (FUN), imidacloprid (IMI) and thiamethoxam (THI) treatments to the control (CON), * indicates P<0.05, ** indicates P<0.01, *** indicates P<0.001 and N.S. indicates not significant.

| Crop | Guild | Sample Type | Taxon | Treatment Mean ± S.E. | | | |
| --- | --- | --- | --- | --- | --- | --- | --- |
|  |  |  |  | CON | FUN | IMI | THI |
| 2015 Full Season Soybean | Predator | VC | Phlaeothripidae | 5.5±1.3 | 5.3±1.4<sup>N.S.</sup> | 2.9±1.1* | 2.3±0.9* |
| | Pest | VC | Cicadellidae | 3.8±1.5 | 4.3±0.8<sup>N.S.</sup> | 2.6±0.9<sup>N.S.</sup> | 0.5±0.3* |
| | | | Thripidae | 13.0±4.0 | 13.4±2.7<sup>N.S.</sup> | 7.5±1.8<sup>N.S.</sup> | 4.3±0.8* |
| 2015-2016 Winter Wheat | Predator | PT | Mesostigmata | 5.1±1.3 | 5.0±1.2<sup>N.S.</sup> | 8.0±1.8* | 5.9±1.6<sup>N.S.</sup> |
| | | LE | Staphylinidae | 2.3±0.5 | 2.4±0.6<sup>N.S.</sup> | 1.0±0.2* | 1.3±0.2<sup>N.S.</sup> |
| | Parasitoid | SC | Aphelinidae | 6.1±1.8 | 5.7±1.3<sup>N.S.</sup> | 2.3±0.6* | 2.7±0.6<sup>N.S.</sup> |
| | | | Braconidae | 2.3±0.4 | 1.9±0.4<sup>N.S.</sup> | 1.4±0.3** | 2.0±0.3<sup>N.S.</sup> |
| | Pest | SC | Thripidae | 20.8±6.4 | 17.0±5.3<sup>N.S.</sup> | 14.1±4.5** | 18.3±8.5* |
| | | VC Fall | Aphididae | 5.9±1.2 | 8.9±2.7 | 1.0±0.5*** | 0.8±0.3*** |
| | | VC Spring | *Oulema melanopus* | 1.8±0.5 | 2.4±0.6<sup>N.S.</sup> | 3.2±0.7** | 2.4±0.9<sup>N.S.</sup> |
| | Other | PT | Acari (Oribatida + Tarsonemidae) | 2.8±0.7 | 4.1±0.8<sup>N.S.</sup> | 7.0±1.8** | 4.3±0.9<sup>N.S.</sup> |
| 2016 Double Cropped Soybean | Predator | LE | Staphylinidae | 0.9±0.2 | 1.7±0.3** | 1.2±0.3<sup>N.S.</sup> | 0.9±0.2<sup>N.S.</sup> |
| | | SN | Araneae | 5.5±1.4 | 3.5±0.4<sup>N.S.</sup> | 2.8±0.9* | 2.4±0.5** |
| | | | Anthocoridae | 4.4±1.1 | 3.9±1.1<sup>N.S.</sup> | 1.1±0.4* | 0.8±0.4** |
| | | | Coccinellidae | 1.4±0.7 | 1.4±0.5<sup>N.S.</sup> | 0.1±0.1* | 0.4±0.3* |
| | | VC | Phlaeothripidae | 7.1±1.5 | 3.9±0.5<sup>N.S.</sup> | 5.0±0.9<sup>N.S.</sup> | 3.9±1.6** |
| | Parasitoid | PT | Scelionidae | 3.5±0.5 | 3.0±0.5<sup>N.S.</sup> | 2.1±0.4* | 3.0±0.5<sup>N.S.</sup> |
| | | SN | Braconidae | 1.8±0.5 | 0.9±0.5<sup>N.S.</sup> | 0.6±0.5** | 1.5±0.7<sup>N.S.</sup> |
| | Pest | SC | Thripidae | 14.6±3.0 | 9.7±2.2* | 14.1±3.4<sup>N.S.</sup> | 11.9±3.8<sup>N.S.</sup> |
| | Other | LE | Collembola | 53.3±12.0 | 78.8±15.3<sup>N.S.</sup> | 87.6±17.0* | 75.2±13.3<sup>N.S.</sup> |
| | | SC | Chloropidae<br>Sciaridae | 58.3±10.5<br>4.7±1.7 | 40.1±7.7*<br>2.4±0.8<sup>N.S.</sup> | 39.0±7.4**<br>1.9±0.5* | 52.2±12.4<sup>N.S.</sup><br>3.5±0.9<sup>N.S.</sup> |
| 2017 Maize | Predator | PT | Araneae | 7.0±1.8 | 4.4±1.0<sup>N.S.</sup> | 3.1±0.5** | 3.0±0.4** |
| | | VC July | Araneae | 8.3±1.6 | 11.9±2.2<sup>N.S.</sup> | 13.8±1.6** | 14.5±1.9*** |
| | | | Chrysopidae | 11.0±1.6 | 9.0±1.5** | 8.6±2.2<sup>N.S.</sup> | 10.9±2.5<sup>N.S.</sup> |
| | Pest | SC | Alticini | 2.5±0.7 | 1.4±0.5* | 0.9±0.3** | 0.6±0.1** |
| | Other | PT | Acari | 7.8±2.1 | 13.9±6.1<sup>N.S.</sup> | 16.0±4.9** | 11.6±4.6<sup>N.S.</sup> |
| | | SC | Phalacridae | 1.9±0.4 | 1.6±0.3<sup>N.S.</sup> | 1.2±0.3* | 1.2±0.3* |

##### 2015-2016 Winter wheat

Imidacloprid treated plots exhibited higher pitfall trap abundances of free-living predatory mites (Mesostigmata, F_3,77_=2.4, P=0.074) and saprovorous mites (Acari - Orbatida +Tarsonemidae, F_3,77_=3.23, P=0.027) (Table 2). However, rove beetles (Staphylinidae, F_3,77_=3.03, P=0.035) were less abundant in the imidacloprid treatment litter extractions than the control, and sticky cards in imidacloprid treated plots captured fewer aphelinid (F_3,77_=4.27, P=0.008) and braconid wasps (F_3,77_=2.98, P=0.037). Fewer phytophagous thrips (Thripidae, F_3,77_=3.18, P=0.029) were captured on sticky cards in both insecticide treatments, and visual scouting indicated that aphids (Aphididae, F_3,46_=16.45, P<0.001) were controlled by both treatments in the fall/winter. However, cereal leaf beetle (*Oulema melanopus*, F_3,77_=5.47, P=0.002) abundances were higher in imidacloprid treated plots compared to the control in the spring.

##### 2016 Double-cropped soybean

In litter extractions, rove beetles (Staphylinidae, F_3,77_=3.29, P=0.025) were more abundant in the fungicide treatment compared to the control, and more collembola (F_3,77_=2.29, P=0.085) were extracted from the litter in the imidacloprid treatment compared to the control (Table 2). Fewer scelionid (F_3,77_=2.33 P=0.081) parasitoid wasps were pitfall trapped and fewer braconid wasps (F_3,15_=4.08, P=0.026) were captured in sweep net samples within imidacloprid treated plots. Visual scouting detected fewer predatory thrips (Phlaeothripidae, F_3,46_=3.04, P=0.038) within thiamethoxam treated plots. Spiders (Araneae, F_3,15_=4.09, P=0.026), minute pirate bugs (Anthocoridae, F_3,77_=5.58, P=0.009), and lady beetles (Coccinelidae, F_3,15_=4.24, P=0.023) were suppressed in sweep net samples from both insecticide treatments. Phytophagous thrips (Thripidae, F_3,77_=3.36, P=0.024) sticky card captures were reduced in the fungicide treatment. Dark-winged fungus gnat (Sciaridae, F_3,61_=2.54, P=0.064 sticky card captures were lower in the imidacloprid treatment, with fewer grass flies (Chloropidae, F_3,61_=3.54, P=0.020) in the fungicide and imidacloprid treatments.

##### 2017 Maize

Similar to winter wheat, more saprovorous mites (Acari - Oribatida +Tarsonemidae, F_3,77_=2.94, P=0.038) were captured in imidacloprid treated pitfall traps (Table 2). Higher abundances of spiders (Araneae, F_3,15_=16.79, P<0.001) were detected with visual sampling in both insecticide treated plots, in contrast to the fewer spiders (F_3,77_=3.55, P=0.018) that were captured in pitfall traps. Green lacewings (Chrysopidae, F_3,15_=3.69, P=0.036) were less abundant in fungicide visual samples. All pesticide treatments captured fewer flea beetles (Chrysomelidae - Alticini, F_3,77_=4.64, P=0.005) on sticky cards than the control, and fewer shining flower beetles (Phalacridae, F_3,77_=2.84, P=0.040) were collected in both insecticide treatments.

### 3.3 Crop sampling

To evaluate treatment impacts on plant growth rates and health, plant height, stand count and yield were measured in all the crops, with NDVI and the number of tillers also measured in wheat (Table 3). Stand count was improved in imidacloprid treated plots compared to the control in full-season soybean (F_3,15_=15.11, P<0.001) and in both insecticide treatments in maize (F_3,15_=6.65, P=0.005). The plant height was also greater in all three pesticide treatments compared to the control in maize (F_3,15_=4.18, P=0.011). Plant height, NDVI and tiller counts in wheat were not impacted by the treatments, but stand count was lower in the thiamethoxam treatment than the control (F_3,46_=2.32, P=0.087). In wheat, the fungicide and imidacloprid treatments increased yield in comparison to the control (F_3,15=_5.16, P=0.012), but yield benefits were not observed in any other crop.

**Table 3.** The effect of seed treatments on plant health parameters and yield for each crop. Analysis of variance was used with treatment as a fixed effect and location, row (location) and column (location) as random effects. † and ^ represent cases with two and four sampling dates, respectively. In these cases, treatment and date were used as fixed effects. For effect differences of P<0.10, contrasts were used to compare the fungicide (FUN), imidacloprid (IMI) and thiamethoxam (THI) treatments to the control (CON). * indicates P<0.05, ** indicates P<0.01, *** indicates P<0.001 and N.S. indicates not significant. Results where contrasts were performed are bolded.

| Metric | Treatment Mean ± S.E. |  |  |  | D.F | Treatment F-value, P-value |
| --- | --- | --- | --- | --- | --- | --- |
|  | CON | FUN | IMI | THI |  |  |
| 2015 Full-season Soybean |  |  |  |  |  |  |
| Stand Count (plants 2m<sup>-1</sup>) | 15.5±1.4 | 16.8±1.4 <sup>N.S.</sup> | 20.2±1.6*** | 15.3±1.6 <sup>N.S.</sup> | 3,15 | 15.11, <0.001 |
| Plant Height (cm) | 26.4±1.0 | 26.7±0.9 | 27.8±1.2 | 27.6±0.8 | 3,15 | 0.78, 0.524 |
| Yield (kg ha<sup>-1</sup>) | 3901±212 | 4113±198 | 4129±180 | 3974±210 | 3,15 | 0.63, 0.609 |
| 2015 – 2016 Winter Wheat |  |  |  |  |  |  |
| Stand Count (plants m<sup>-1</sup>) † | 42.0±2.6 | 40.0±3.4 <sup>N.S.</sup> | 42.0±2.5 <sup>N.S.</sup> | 38.3±2.5* | 3,46 | 2.32, 0.087 |
| Plant Height (cm) | 13.5±0.4 | 13.0±0.2 | 13.5±0.3 | 13.1±0.3 | 3,15 | 1.42, 0.276 |
| No. of Tillers (tillers 0.9m<sup>-1</sup>) † | 137±4 | 147±8 | 132±6 | 134±7 | 3,46 | 1.33, 0.275 |
| NDVI ^ | 0.40±0.01 | 0.40±0.01 | 0.40±0.01 | 0.40±0.01 | 3,108 | 0.06, 0.981 |
| Yield (kg ha<sup>-1</sup>) | 2845±277 | 3373±201* | 3584±213** | 3383±389 <sup>N.S.</sup> | 3,15 | 5.16, 0.012 |
| 2016 Double-cropped Soybean |  |  |  |  |  |  |
| Stand Count (plants 2m<sup>-1</sup>) † | 16.6±0.7 | 17.9±0.7 | 16.7±0.7 | 18.0±0.6 | 3,46 | 2.01, 0.126 |
| Plant Height (cm) | 52.0±3.0 | 56.4±1.6 | 55.4±3.0 | 54.6±3.6 | 3,15 | 1.22, 0.336 |
| Yield (kg ha<sup>-1</sup>) | 3068±184 | 3165±179 | 3203±169 | 3148±178 | 3,15 | 0.31, 0.821 |
| 2017 Maize |  |  |  |  |  |  |
| Stand Count (plants 2m<sup>-1</sup>) | 11.6±0.5 | 11.8±0.4 <sup>N.S.</sup> | 12.2±0.4** | 12.0±0.5** | 3,15 | 6.65, 0.005 |
| Plant Height (cm)† | 29.4±4.0 | 31.2±4.0* | 31.9±4.2** | 31.5 ± 4.2* | 3,46 | 4.18, 0.011 |
| Yield (kg ha<sup>-1</sup>) | 8850±485 | 9595±204 | 9583±711 | 9502±520 | 3,15 | 0.58, 0.640 |

## 4. Discussion

We conducted a three-year field study evaluating pesticide seed treatment impacts in a full-season soybean, winter wheat, double-cropped soybean and maize rotation. Our specific goals were to quantify neonicotinoid residues in the soil and in winter annual flowers, which underlies the magnitude of non-target impacts on the arthropod community. We also quantified benefits to plant growth and yield to determine whether treatments were economically justified. Trace amounts of insecticide were present in one winter annual species in one site year, which did not correspond with our treatments. Low levels of insecticide residues were present in the soil, with the highest levels observed in the final year, suggesting some accumulation. Pesticide seed treatments variably impacted the arthropod community throughout the study. Shannon diversity indices, taxa evenness, and PRC analyses demonstrated occasional community disturbances of a relatively small magnitude, and pesticide seed treatments also impacted individual taxa. However, there was little consistency between crops and sampling methods. Overall, the imidacloprid treatment had stronger impacts than the thiamethoxam treatment, and the fungicide only treatment also occasionally impacted arthropod communities. Pest pressure was very low throughout the study, and while the treatments occasionally improved early season plant growth, we only observed yield benefits in winter wheat, which we attributed to the fungicide component since it occurred across pesticide treatments.

### 4.1 Environmental persistence and routes of exposure to neonicotinoid residues

#### 4.1.1 Persistence in plants

Neonicotinoid insecticides are systemic (Nauen et al., 2008), with active ingredient taken up by crop plants and distributed throughout their tissues (Alford & Krupke, 2017; Myers & Hill, 2014). In maize and soybean, neonicotinoid seed treatment active ingredients only remain active in plant tissue for three to four weeks post planting (Alford & Krupke, 2017; Myers & Hill, 2014). In fall planted winter wheat, the exact activity period is unknown; Zhang et al. (2016) found trace amounts of imidacloprid and clothianidin in seed treated winter wheat up to 200 days after planting and observed successful control of cereal aphids throughout the growing period. The presence of insecticide in plant tissue over a longer period could be a source of exposure for non-target beneficials arthropods such as lady beetles and minute pirate bugs that supplement their diet with plant material, or parasitoids that rely on nectar as a food source (Gontijo et al., 2015; Moscardini et al., 2014; Moser & Obrycki, 2009). Neonicotinoids could remain active for much longer in winter wheat than in maize and soybean, because of low temperatures and plant dormancy during the winter and early spring.

Neonicotinoid residues can also be taken up from the soil by non-target plants, such as wildflowers and inter-seeded cover crops (Botías et al., 2015; Bredeson & Lundgren, 2019; Krupke, Hunt, Eitzer, Andino, & Given, 2012; Pecenka & Lundgren, 2015); these are important resources for pollinators, and could be another source of neonicotinoid exposure (Bretagnolle & Gaba, 2015; Mandelik, Winfree, Neeson, & Kremen, 2016). Since these non-target plants were sampled during peak planting and crop production seasons, aerial deposition cannot be separated from uptake. To mitigate this issue, we sampled in late winter. Trace levels of imidacloprid were present in *S. media* flower samples at Beltsville in 2017. Neonicotinoid levels were below the detection threshold for our analysis (5 ppb) and did not correspond with our treatments. Previous studies quantifying residues within non-target plants often detected levels of less than 5 ppb (Bredeson & Lundgren, 2019; Pecenka & Lundgren, 2015); therefore, despite low soil residues, winter annual flowers may uptake small amounts of active ingredient.

#### 4.1.2 Persistence in soil

In soil, the half-life of neonicotinoids can vary greatly, ranging from 28-1250 days for imidacloprid, and 7-353 days for thiamethoxam (Goulson, 2013), with temperature, sunlight, and soil texture, organic matter and moisture content impacting persistence (Bonmatin et al., 2015). Persistence in soil also varies by the amount of active ingredient used, which can differ greatly between crops due to different treatment and seeding rates. We did not detect high levels of neonicotinoid residues in the soil, but the highest levels of both insecticides were observed after 2017 maize planting, suggesting the possibility of some accumulation across crops, as hypothesized. This was further supported by higher imidacloprid levels in imidacloprid treated plots than surrounding plots prior to 2017 maize planting at Beltsville. Overall, imidacloprid was detected more often than thiamethoxam, with some detected before the start of the study at Beltsville, even though imidacloprid was not used in that field the previous year. This difference in soil persistence is likely due to imidacloprid’s longer half-life. High moisture content, temperature and sunlight are all positively correlated with neonicotinoid breakdown, and thiamethoxam and imidacloprid also have high leaching potential (Bonmatin et al., 2015). Given the high summer temperatures and precipitation in Maryland, the low levels of neonicotinoid residues in our plots could be caused by rapid microbial and photolytic breakdown of residues, or by leaching and run-off.

### 4.2 Non-target impacts of NSTs on arthropods

#### 4.2.1 Impacts on communities over time

The arthropod community was impacted variably in different crops and sampling types. Pitfall trap data did not show community disturbance through PRC analysis, diversity, or evenness metrics. Litter communities exhibited lower diversity in both insecticide treatments and lower evenness in the imidacloprid treatment before planting maize in 2017, as well as lower diversity in the imidacloprid treatment and lower evenness in all three pesticide treatments after planting maize. Our hypothesis that the soil community would experience the strongest impacts was not supported, as we generally observed a greater impact on foliar taxa. However, the results for the litter community provide some support for the hypothesis that the soil community would experience greater disturbance later in the study due to accumulation of active ingredients in the soil and litter. Peck (2009) used pitfall traps and soil core extractions to measure the impact of repeated imidacloprid applications in turf, and found that only taxa extracted from soil cores were impacted, with reduced abundance of collembola, true bugs, ground beetles and rove beetles. Conversely, Disque et al. (2018) observed deviation from the control in PRC analysis of pitfall data but not litter data from clothianidin treated maize. While litter or soil core extractions are absolute samples of the community at a given time and place, pitfall traps measure activity density; therefore, these sampling methods may vary in their effectiveness in capturing treatment impacts.

Sticky card, sweep net, and visual sample data exhibited pesticide treatment impacts on the arthropod community. The PRC analysis for sticky cards showed increasing disturbance over time in both winter wheat and maize, with no recovery over the sampling period. Given the short period of neonicotinoid activity in crop plants, we expected sticky card and foliar communities to recover, as observed by Disque et al. (2018) in maize; our results suggest that NSTs may have longer term effects on sticky card arthropod communities than previously observed. Redundancy analysis of sweep net data demonstrated community disturbances in 2016 double-cropped soybean but not 2015 full-season soybean. In 2016, diversity was lower in the thiamethoxam treatment than the control. Taxon diversity and community evenness are both correlated with lowered pest pressure (Lundgren & Fausti, 2015); by reducing diversity and evenness, NSTs could interfere with natural pest control. We did not observe increases in pest abundances that would suggest a breakdown of natural pest control, except for Cereal leaf beetle *Oulema melanopus* in imidacloprid treated wheat, which did not correspond with reductions in diversity and evenness. Instead, treatments may have impacted control of Cereal leaf beetle by natural enemies. For example, neonicotinoids could reduce the presence of alternate prey for natural enemies, preventing their population from building up sufficiently to control Cereal leaf beetle (Yoo & O’Neil, 2009), or exposure to neonicotinoids could reduce the ability of parasitoid wasps to parasitize hosts (Rogers & Potter, 2003).

#### 4.2.2. Impacts on individual taxa within a crop

Through the different crops and sampling methods, we observed impacts on taxa that fill various ecological roles in the agricultural arthropod community. The soil community was dominated by mites and collembola, which demonstrated numerically higher abundances in the pesticide treatments throughout the study. Previous studies have found that neonicotinoids can have both positive and negative impacts on collembola (Peck, 2009; Zaller et al., 2016) and can stimulate fecundity in mites (Pisa et al., 2015). Disque et al. (2018) also observed increased collembola and mite activity density in a similar study conducted with clothianidin treated maize, which could be attributed to a trophic cascade caused by natural enemy suppression, or changes in the fungal community caused by the fungicide components. Both soil mites and collembola play a key role in breaking down soil organic matter (Crossley, Mueller, & Perdue, 1992), and by altering the abundance or activity of these organisms, NSTs could further disturb agroecosystems.

In pitfall and litter samples, we observed suppression of two prominent groups of generalist predators in insecticide treated crops, rove beetles and spiders. Spider abundance was also reduced in both insecticide treatments in double-cropped soybean sweep net samples. The low toxicity of neonicotinoids towards arachnids (Douglas & Tooker, 2016) suggests that changes in spider density were driven by prey scarcity or other community interactions. In contrast, spider abundance was higher in the maize visual samples from the insecticide treated plots. These contradictory results could be due to variation in behavioral responses to the active ingredients; Easton and Goulson (2013) found that spiders were attracted to low doses of imidacloprid, but were repelled by a high dose. In sweep net and visual samples, we observed reduced abundance or activity density of various predators that have previously been shown to be impacted by NSTs, such as minute pirate bugs (Anthocoridae), lady beetles (Coccinellidae), and predatory thrips (Phlaeothripidae) (Albajes, López, & Pons, 2003; Amjad, Azam, Sarwar, Malik, & Sattar, 2018; Disque et al., 2018; Gontijo et al., 2015; Seagraves & Lundgren, 2012; Zhang et al., 2016). The imidacloprid treatment also suppressed aphelinid and braconid wasps captured on sticky cards in winter wheat, and braconid wasps in double cropped soybean sweep net samples. Parasitoids are especially important for controlling cereal aphids in wheat (Schmidt et al., 2003), therefore suppression of aphid natural enemies could have economic implications.

#### 4.2.3 Potential for sublethal impacts

While we only measured abundance or activity density, neonicotinoids can negatively impact behavior, condition, reproductive success and survival of non-target arthropods (Main, Webb, Goyne, & Mengel, 2018). Examples of sublethal impacts on predatory taxa include reduced survival, longevity and oviposition in lady beetles (Papachristos & Milonas, 2008), delayed development and reduced fecundity and survival in minute pirate bugs (Gontijo et al., 2015), and paralysis, impaired walking, and increased grooming in ground beetles (Kunkel, Held, & Potter, 2001). In parasitoids, exposure to sublethal doses of neonicotinoids can decrease longevity, disrupt courtship behavior, impair mate and host-finding ability, as well as reduce parasitism rates and the ratio of female offspring (Frewin, Schaafsma, & Hallett, 2014; Moscardini et al., 2014; Stapel, Cortesero, & Lewis, 2000; Tappert, Pokorny, Hofferberth, & Ruther, 2017). Such sublethal effects may have contributed to the patterns observed in our study. The increasing disruption of the community over time that occurred in wheat and maize sticky cards could be due to the pesticides reducing reproductive success, which would emerge over multiple generations.

#### 4.2.3 Differences between pesticides

In addition to impacts due to both insecticide treatments, there were several instances where only the imidacloprid treatment exhibited impacts. For example, thiamethoxam did not affect parasitoids, while imidacloprid reduced braconid, aphelinid and scelionid wasps. Burgess and King (2015) previously found that the LC_50_ of imidacloprid was 4.5 times lower than that of thiamethoxam for a parasitoid of houseflies. Imidacloprid has a longer half-life and is an older chemical than thiamethoxam (Bonmatin et al., 2015; Simon-Delso et al., 2015), which could explain its greater non-target impacts.

We included a fungicide only treatment in order to isolate effects of seed applied fungicides, and the results supported our hypothesis that these fungicides would impact the arthropod community. For example, in double-cropped soybean, the fungicide only treatment solely reduced phytophagous thrips abundance on sticky cards and increased abundance of rove beetles in litter samples. Additionally, the fungicide only treatment exhibited similar impacts to one or both insecticide + fungicide treatments in certain cases, such as similar deviations from the control in several PRC analyses. To the best of our knowledge, few studies have evaluated the persistence of seed applied fungicides in agroecosystems, or their impact on the arthropod community, even though they can be moderately toxic to arthropods (MDA, 2012). Both fungicide and insecticide seed treatments decreased earthworm surface activity and increased collembola surface activity in wheat (Van Hoesel et al., 2017; Zaller et al., 2016). Given that the fungicide treatments consist of several active ingredients, those ingredients could interact synergistically with each other or with the insecticides to impact the arthropod community. The effects of fungicides on arthropod health have been investigated in pollinators; clothianidin can synergistically interact with the fungicide propiconazole leading to higher mortality in multiple bee species (Sgolastra et al., 2017). In addition, fungicides could alter arthropod abundance by interfering with entomopathogenic fungi, thereby altering disease pressure (Lagnaoui & Radcliffe, 2009). In our study, the soil community was dominated by fungivore taxa (mites and collembola). Therefore, fungicides could also affect arthropods through changes in fungal diversity and abundance, impacting resources available for fungivores. Regardless of the mechanism, our results clearly demonstrate that seed applied fungicides can disrupt arthropod communities in agroecosystems.

### 4.3 Economic impacts

Although NSTs suppressed pests throughout the study, namely thrips (Thripidae) and leafhoppers (Cicadelldiae) in soybean, aphids (Aphididae) in early season wheat, and flea beetles (Chrysomelidae – Alticini) in maize, these pests were not present at economically damaging levels. As we predicted, the insecticides did not improve yield through pest suppression. The insecticide treatments improved early season stand density and plant height, but these benefits did not translate to yield increases. In wheat, the fungicide and imidacloprid treatments significantly increased yield, which was also numerically higher in the thiamethoxam treatment. Given that yield improvements occurred across the pesticide treatments, it was likely a result of the fungicide products and could be achieved without insecticides. Our results are consistent with several previous findings that NSTs may not provide economic benefits in the absence of early season pest pressure (Cox et al., 2007; Myers & Hill, 2014; Wilde et al., 2007). This suggests that the use of NSTs in Maryland grain production may not be warranted outside of specific instances of high pest pressure.

## 5. Conclusions and Future Directions

We found that NSTs can disturb arthropod communities in Maryland grain systems, despite low levels of neonicotinoid residues in the agroecosystem, and the communities occasionally were unable to recover by the end of the sampling period. While we observed treatment impacts on diversity metrics as well as individual taxa abundances, results were inconsistent between crops and sampling methods, making it difficult to elucidate the underlying mechanisms. The persistence of neonicotinoids in the environment can vary greatly, and the effects on the community in our study suggest that areas with greater neonicotinoid persistence and accumulation could experience much stronger disturbances. This study is among the first to document disruption of arthropod communities by seed applied fungicides that are used with NSTs, which have received little attention with regards to their impacts on arthropods. Given their inclusion in all commercial neonicotinoid seed treatments, there is an urgent need to further understand their role. Without a corresponding increase in pest pressure (Douglas & Tooker, 2015), NST treated maize and soybean acreage has increased, with many of these acres not previously treated with insecticides. Between 2011 and 2014, the overall quantity of neonicotinoids applied to maize also doubled, indicating an increase in the rate of products used (Tooker, Douglas, & Krupke, 2017). Despite minimal or no benefits in many cases, NST use has continued to grow. Unfortunately, there is little availability of maize without NSTs in the US, leaving farmers with limited choices (Alford & Krupke, 2017). Given the levels of NST contamination in the environment and the impacts on non-target arthropod communities, tactics must be developed to minimize overuse.

## 6. Author Contributions

A.D., G.D., and K.H. developed and carried out the study, conducted data analysis and wrote the manuscript. M.L. participated in data collection and manuscript preparation.

## 7. Acknowledgements

This project was supported by the Hatch/Multistate funds [project no. MD-ENTM-8887/project accession no. 1009567 and project no. MD-ENTO-9589/project accession no. 1012455] from the USDA National Institute of Food and Agriculture, by USDA NIFA award number 2015-38640-23777 through the North East SARE program under sub-award number GNE16-11B-29994, and by the Maryland Grain Producers Utilization Board and the Maryland Soybean Board.

Outreach efforts and publication costs associated with this project were supported by the Crop Protection and Pest Management Program [grant no. 2017-70006-27171/project accession no. 1013913] from the USDA National Institute of Food and Agriculture. Opinions, findings, conclusions, or recommendations expressed in this publication are those of the author(s) and do not necessarily reflect the view of these organizations.

We would like to thank Kevin Conover, John Draper, and the staff at the Central Maryland and Wye Research and Education Centers for planting and maintaining our study plots, and Terry Patton, Matt Dimmock, Robert Starkenburg and the other members of the Hamby and Dively Labs who contributed to this project. We would also like to thank Stephanie Yarwood, Daniel Gruner, and Robert Kratochvil for their constructive reviews.

## 8. Data Accessibility

Data will be made available through the Dryad Digital Repository.

