## Supporting Information Document for "Evaluating the ecological impacts of pesticide seed treatments on arthropod communities in a grain crop rotation"

### 1. Methods

For all arthropod and crop sampling, the outer 1m of the plot was excluded on all sides to avoid edge effects.

#### *1.1 Residue analysis*

##### *1.1.1 Winter annual flower collection*

In 2016, flower buds were collected at both sites on March 15. To collect sufficient material for analysis, samples from two replicates of the same treatment (Column 1+ Column 2; Column 3+ Column 4) (Fig. S1) were combined at both sites. In 2017, common henbit was collected at Beltsville on March 13 and common chickweed at Queenstown on April 12, with samples combined as described previously. At Beltsville, we also collected common chickweed on April 10, and were able to collect enough material for analysis from each individual plot. Flower buds (3g) were stored at -80°C in falcon tubes until they were sent for neonicotinoid residue analysis.

##### *1.1.2 Soil collection*

We took 30 random soil cores (1.9cm diameter and 12cm depth) from within and between rows in each plot and mixed them into a single homogenized sample. Due to space restrictions, only a small subsample of each homogenized sample was collected and stored at -80ºC for subsequent analysis. Because these samples were also used for another experiment, insufficient soil was available to analyze residues within each plot individually. In maize and soybean, soil from two replicates of the same treatment (Col 1+ Col 2; Col 3+ Col 4) (Fig. S1) were combined, and in wheat, soil from all four replicates was combined.

#### *1.1.3 Analysis*

Winter annual flower buds and soil were sent to the USDA National Science Laboratory (Gastonia, North Carolina, USA) for analysis. Briefly, neonicotinoid residues were extracted with a refined official pesticide extraction method [AOAC OMA 2007.0, the QuEChERS method (Quick, Easy, Cheap, Effective, Rugged, and Safe)], using an acetonitrile and water solution. Extraction was followed by enhance matrix reduction (EMR) clean-up and analysis using certified standard reference materials and liquid chromatography coupled with tandem mass spectrometry detection (LC/MS/MS) utilizing the precursor and product ions of analytes of interest. The detection level was 10 ppb for imidacloprid, 5 ppb for thiamethoxam and 30 ppb for clothianidin in flowers, and in soil was 5ppb for imidacloprid, 10ppb for thiamethoxam and 15ppb for clothianidin in soil.

### *1.2 Arthropod sampling*

#### *1.2.1 Pitfall traps*

Pitfall traps consisted of two stacked 360ml plastic cups buried so that the opening was level with the soil surface. The inner cup contained approximately 60ml of ethylene glycol and was sheltered from weather and wildlife interference with a 30cm square black plastic cover supported by three carriage bolts held approximately 5cm above the soil surface. In 2015 and 2016, we used 50% ethylene glycol, but switched to 100% in 2017 due to the combination of diluted ethylene glycol and water from rainfall allowing samples to degrade. Three pitfall traps were set up between the rows within each plot in an evenly spaced diagonal line. On each sampling date, pitfall traps were left in place for one week. After collection, samples were vacuum filtered and transferred into alcohol that was dyed with food coloring to increase visibility of soft bodied arthropods. Data from the three pitfall subsamples per plot were averaged before analysis.

*1.2.2 Litter extraction*

Litter samples were collected by using masonry trowels to gather all the litter and the soil rhizosphere to a depth of about 1cm from a circular (full season soybean and maize) or rectangular (double cropped soybean and wheat) $0.09m^2$ area. Four sets of litter were collected per plot, and the litter was combined into two subsamples for extraction with Berlese funnels. Burlese funnels were constructed using a 19L painter's bucket with the bottom replaced by a layer of wide mesh, placed over a metal funnel with a cup of 70% ethanol placed underneath. The alcohol was dyed with food coloring to aid in visualization of soft bodied arthropods. 30 W incandescent bulbs were used as the heat source during extraction, and samples were placed in the funnel for up to 48 hours, until the soil was completely dry. The subsamples were averaged before analysis.

*1.2.3 Sticky cards*

One sided yellow sticky cards (7.62 cm x 12.7 cm) were placed horizontally at a height of 8 cm. Three sticky cards were placed in an evenly spaced diagonal line between rows in each plot. On each sampling date, sticky cards were deployed for one week. Data from the three sticky cards subsamples was averaged before analysis.

*1.2.4 Visual inspection*

In soybean, leaf trifoliate inspections were conducted on the newest trifoliate from 10 randomly selected plants (V5 stage in 2015, V2 and R1 in 2016). For soybean analyses, data from the 10 plants was summed because of low arthropod numbers. In wheat, we scouted for pests twice in the winter and three times in the spring and summer. Two subsamples of two linear yards (1.8 m) were scouted per plot by throwing out a yardstick and counting pests on the plants on either side. Data from the wheat subsamples was averaged for analysis. Earlier in the season, the whole plant was visually examined, but in the last set of samples only the flag leaf and head were included. In maize, foliar arthropods were recorded through visual counts at three time points. At V7, five adjacent plants were scouted from four subsamples taken at randomly selected points, and the whole plant was examined. Data from the subsamples was averaged for analysis. At R1, 10 plants were randomly selected from the middle six rows and were destructively sampled, with all the leaves stripped off the plant and examined. Insects that may have been present within the stem were not included. At R3-R4, plants were selected the same way as R1, but only the ears were sampled. For corn analyses, the sum of data from the all plants was used, due to low arthropod abundance.

*1.3 Crop sampling*

*1.3.1 Stand density*

In all crops, stand density was measured by throwing out a meter stick at randomly selected points and counting the number of plants along it, and data from subsamples was averaged for analysis. In soybean, stand density was measured at emergence in 2015, and at emergence and one-week post emergence in 2016. The meter stick was thrown twice and the number of plants on either side was counted. In wheat, stand density was measured at emergence and one-week post emergence. The meter stick was thrown four times and the number of seedlings on one side of the stick was counted. In maize, stand density was measured shortly after emergence with four throws where the number of plants on both sides of the stick was counted.

*1.3.2 Plant height*

In soybean, plant height was measured concurrently with trifoliate sampling, using the same ten plants per plot, extended to their full height. Data from the 10 plants was averaged for analysis. In wheat, plant height was measured six weeks post planting by randomly throwing out meter sticks at four points and measuring the heights of five randomly selected plants along the meter stick. The heights of the five plants in each subsample were averaged, and then the data from the four subsamples was averaged again. In maize, height was measured along with stand density and again at V7 by measuring the first five plants on one side of the meter stick while fully extended. The height from the five plants was averaged, and then the mean heights from the four subsamples were averaged again.

*1.3.3 Tiller counts*

In wheat, three one-foot (30.4 cm) sections of plants were selected randomly and dug up at Feekes stages 2 and 6. The plants from the three sections were combined and brought back to the lab to count the number of tillers per three row feet.

*1.3.4 Normalized difference vegetation index (NDVI)*

NDVI is a measure of photosynthetic activity that is calculated using the variation in reflectance of light by different surfaces. These measurements were taken three times in the winter and once in the spring using a Crop Circle optical sensor (Holland Scientific, Lincoln, Nebraska, USA). Measurements were taken by walking the length of each plot between the two center rows, while holding the sensor out at shoulder height.

*1.3.5 Yield*

Yield was measured directly by the combine harvester in all cases except wheat and double cropped soybean at Beltsville, where the harvested grain was transferred to a weigh wagon. In full season soybean, the whole plot was harvested at both sites through three passes of a 10-foot combine harvester. In wheat, half of each plot was harvested through three passes of a 5-foot combine harvester through the center of the plot. The wheat yield data from Beltsville was more variable due to a horseweed *Erigeron canadensis* outbreak throughout the field. In double cropped soybean, whole plots were harvested at Beltsville (three 10-foot passes) and half of each plot was harvested at Queenstown (three 5-foot passes). In maize, half of each plot was harvested at both sites (three 5-foot passes). The data was corrected to the appropriate moisture content for each crop and converted to kilograms per hectare for analysis.

### 2. Supporting Tables and Figures

Table S1. Seed treatment active ingredients used in 2015 full-season (FS) soybean and 2015-2016 winter wheat. Soybean variety P93Y84 (Pioneer) was treated at a low rate, which is the most commonly used in Maryland, with a seeding rate of 155,000 seeds per acre at Beltsville (BV) and 150,000 seeds at Queenstown (QT). For wheat, variety MBX14K297 (Mercer) was treated at a medium rate, which was chosen because NSTs are not widely used in Maryland wheat. The same seeding rate was used at both sites (1.75 million seeds per acre).

| Crop | Treatment | Product | Active ingredient (ai) | mg ai seed$^{-1}$ | mg ai plot$^{-1}$ (BV/QT) | g ai ha$^{-1}$ (BV/QT) |
|---|---|---|---|---|---|---|
| 2015 FS Soybean | Thiamethoxam + Fungicide | Cruiser 5FS | Thiamethoxam | 0.0756 | 400.52; 387.60 | 28.96; 28.02 |
| | | Maxim 4FS | Fludioxonil | 0.0038 | 20.13; 19.48 | 1.46; 1.41 |
| | | Apron XL | Mefenoxam | 0.0113 | 59.87; 57.93 | 4.33; 4.19 |
| | | Vibrance | Sedaxane | 0.0038 | 20.13; 19.48 | 1.46; 1.41 |
| | Imidacloprid + Fungicide | Gaucho 600 | Imidacloprid | 0.1000 | 529.78; 512.69 | 38.30; 37.07 |
| | | Allegiance FL | Metalaxyl | 0.0244 | 129.27; 125.10 | 9.35; 9.04 |
| | | Evergol Energy | Prothioconazole | 0.0081 | 42.91; 41.53 | 3.10; 3.00 |
| | | | Penflufen | 0.0045 | 23.84; 23.07 | 1.72; 1.67 |
| | | | Metalaxyl | 0.0064 | 33.91; 32.81 | 2.45; 2.37 |
| | Fungicide Only | Maxim 4FS | Fludioxonil | 0.0038 | 20.13; 19.48 | 1.46; 1.41 |
| | | Apron XL | Mefenoxam | 0.0113 | 59.87; 57.93 | 4.33; 4.19 |
| | | Vibrance | Sedaxane | 0.0038 | 20.13; 19.48 | 1.46; 1.41 |
| 2015-2016 Winter Wheat | Thiamethoxam + Fungicide | Cruiser 5FS | Thiamethoxam | 0.0143 | 854.56 | 61.78 |
| | | Vibrance Extreme | Sedaxane | 0.0013 | 78.62 | 5.68 |
| | | | Difenoconazole | 0.0063 | 377.35 | 27.28 |
| | | | Mefenoxam | 0.0016 | 94.37 | 6.82 |
| | Imidacloprid + Fungicide | Gaucho 600 | Imidacloprid | 0.0217 | 1297.51 | 93.80 |
| | | Allegiance FL | Metataxyl | 0.0017 | 102.27 | 7.39 |
| | | Evergol Energy | Prothioconazole | 0.0018 | 106.01 | 7.66 |
| | | | Penflufen | 0.0009 | 53.00 | 3.83 |
| | | | Metataxyl | 0.0014 | 84.75 | 6.13 |
| | Fungicide Only | Vibrance Extreme | Sedaxane | 0.0013 | 78.62 | 5.68 |
| | | | Difenoconazole | 0.0063 | 377.35 | 27.28 |
| | | | Mefenoxam | 0.0016 | 94.37 | 6.82 |

Table S2. Seed treatment active ingredients (ai) used in 2016 double-cropped (DC) soybean and 2017 maize. Both crops were treated at a medium rate, which is the most commonly used in Maryland. Soybean variety P39T67R (Pioneer) was treated at a rate of 200,000 seeds per acre at Beltsville (BV) and 123,000 seeds per acre at Queenstown (QT). Maize variety TA506-22SPRIb (T.A. Seeds) was treated at 30,000 seeds per acre at Beltsville and 33,000 seeds per acre at Queenstown.

| Crop | Treatment | Product | Active ingredient (ai) | mg ai seed$^{-1}$ | mg ai plot$^{-1}$ (BV/QT) | g ai ha$^{-1}$ (BV/QT) |
|---|---|---|---|---|---|---|
| 2016 DC Soybean | Thiamethoxam + Fungicide | Cruiser 5FS | Thiamethoxam | 0.0756 | 516.80; 317.83 | 37.36; 22.98 |
| | | Maxim 4FS | Fludioxonil | 0.0038 | 25.98; 15.98 | 1.88; 1.15 |
| | | Apron XL | Mefenoxam | 0.0113 | 77.25; 47.51 | 5.58; 3.43 |
| | | Vibrance | Sedaxane | 0.0038 | 25.98; 15.98 | 1.88; 1.15 |
| | Imidacloprid + Fungicide | Gaucho 600 | Imidacloprid | 0.1000 | 683.59; 420.41 | 49.42; 30.39 |
| | | Allegiance FL | Metalaxyl | 0.0244 | 166.80; 102.58 | 12.06; 7.42 |
| | | Evergol Energy | Prothioconazole | 0.0081 | 55.37; 34.05 | 4.00; 2.46 |
| | | | Penflufen | 0.0045 | 30.76; 18.92 | 2.22; 1.37 |
| | | | Metalaxyl | 0.0064 | 43.75; 26.91 | 3.16; 1.95 |
| | Fungicide Only | Maxim 4FS | Fludioxonil | 0.0038 | 25.98; 15.98 | 1.88; 1.15 |
| | | Apron XL | Mefenoxam | 0.0113 | 77.25; 47.51 | 5.58; 3.43 |
| | | Vibrance | Sedaxane | 0.0038 | 25.98; 15.98 | 1.88; 1.15 |
| 2017 Maize | Thiamethoxam + Fungicide | Cruiser 5FS | Thiamethoxam | 0.5000 | 512.69; 563.96 | 37.07; 40.77 |
| | | Vibrance | Sedaxane | 0.0125 | 12.82; 14.10 | 0.93; 1.02 |
| | | Maxim Quattro | Fludioxonil | 0.0063 | 6.46; 7.11 | 0.47; 0.51 |
| | | | Mefenoxam | 0.0050 | 5.13; 5.64 | 0.37; 0.41 |
| | | | Azoxystrobin | 0.0025 | 2.56; 2.82 | 0.19; 0.20 |
| | | | Thiabendazole | 0.0501 | 51.37; 56.51 | 3.71; 4.09 |
| | Imidacloprid + Fungicide | Gaucho 600 | Imidacloprid | 0.5000 | 512.69; 563.96 | 37.07; 40.77 |
| | | Vortex FL | Ipconazole | 0.0063 | 6.46; 7.10 | 0.47; 0.51 |
| | | Allegiance FL | Metalaxyl | 0.0050 | 5.17; 5.68 | 0.37; 0.41 |
| | | Trilex Flowable | Trifloxystrobin | 0.0126 | 12.91; 14.21 | 0.93; 1.03 |
| | Fungicide Only | Vibrance | Sedaxane | 0.0125 | 12.82; 14.10 | 0.93; 1.02 |
| | | Maxim Quattro | Fludioxonil | 0.0063 | 6.46; 7.11 | 0.47; 0.51 |
| | | | Mefenoxam | 0.0050 | 5.13; 5.64 | 0.37; 0.41 |
| | | | Azoxystrobin | 0.0025 | 2.56; 2.82 | 0.19; 0.20 |
| | | | Thiabendazole | 0.0501 | 51.37; 56.51 | 3.71; 4.09 |

Table S3. Timeline for crop and arthropod sampling in 2015 full season soybean. The two dates represent the sampling date at Beltsville and Queenstown, respectively. Soybean was planted on 5/14 at Beltsville and 5/26 at Queenstown and harvested on 10/22 at both sites.

| **Sample Type** | **Growth Stage** | | | |
|---|---|---|---|---|
| | **Pre-Planting** | **VC-V2** | **V5** | **R3** |
| Stand Count | | 5/28, 6/5 | | |
| Height | | | 6/23, 7/1 | |
| Visual Count | | | 6/23, 7/1 | |
| Sticky Card | | 6/2, 6/12 | 6/24, 7/1 | 8/3, 8/14 |
| Pitfall Trap | 5/12, 5/13 | 6/2, 6/12 | 6/24, 7/1 | |
| Litter Extraction | 5/12, 5/21 | 6/2, 6/12 | 6/26, 7/1 | |
| Sweep Net | | | | 8/7, 8/14 |

Table S4. Timeline for crop and arthropod sampling in 2015–2016 winter wheat. October to December dates are from 2015 while March to June dates are from 2016. The two dates represent the sampling date at Beltsville and Queenstown, respectively. Growth stages were measured using the Feekes scale. Wheat was planted on 10/26 at Beltsville and 10/27 at Queenstown and harvested on 6/30 at Beltsville and 6/29 at Queenstown. Two sets of dates in a single cell indicate that sampling occurred twice during that growth stage.

| **Sample Type** | **Growth Stage** | | | | | |
|---|---|---|---|---|---|---|
| | **Stage 1** | **Stage 2** | **Stage 4** | **Stage 5-6** | **Stage 9-10** | **Stage 11** |
| Stand Count | 11/4, 11/5<br>11/11, 11/11 | | | | | |
| Height | | 12/4, 12/4 | | | | |
| NDVI | 11/11, 11,11 | 12/4, 12/4<br>12/16, 12/16 | 3/2, 3/7 | | | |
| Tiller Count | | 12/16, 12/16 | | 4/15, 4/14 | | |
| Visual Count | | 12/4, 12/4<br>12/16, 12/16 | | 4/15, 4/14 | 5/19, 5/16 | 6/10, 6/7 |
| Sticky Card | | | | 4/28, 4/25 | 5/26, 5/25 | 6/17, 6/14 |
| Pitfall Trap | | | | 4/28, 4/25 | 5/26, 5/25 | 6/17, 6/14 |
| Litter Extraction | | | | 4/28, 4/25 | 5/26, 5/25 | 6/17, 6/14 |

Table S5. Timeline for crop and arthropod sampling in 2016 double cropped soybean. The two dates represent the sampling date at Beltsville and Queenstown, respectively. Soybean was planted on 7/8 and harvested on 11/2 at both sites.

| **Sample Type** | **Growth Stage** | | | |
|---|---|---|---|---|
| | **VE-VC** | **V1-V3** | **R1** | **R3** |
| Stand Count | 7/21, 7/19 | 7/28, 7/26 | | |
| Height | | | 8/25, 8/23 | |
| Visual Count | | 7/28, 7/26 | 8/25, 8/23 | |
| Sticky Card | | 7/28, 7/26 | 8/25, 8/23 | 9/12, 9/13 |
| Pitfall Trap | | 7/28, 7/26 | 8/25, 8/23 | 9/12, 9/13 |
| Litter Extraction | | 7/28, 7/26 | 8/25, 8/23 | 9/12, 9/13 |
| Sweep Net | | | 8/25, 8/23 | |

Table S6. Timeline for crop and arthropod sampling in 2017 maize. The two dates represent the sampling date at Beltsville and Queenstown, respectively. Maize was planted on 5/4 at Beltsville and 5/8 at Queenstown and harvested on 10/5 at Beltsville and 9/27 at Queenstown.

| **Sample Type** | **Growth Stage** | | | | | |
|---|---|---|---|---|---|---|
| | **Pre-planting** | **V3-V4** | **V7** | **V10-V12** | **R1** | **R3** |
| Stand Count | | 5/22, 5/24 | | | | |
| Height | | 5/22, 5/24 | 6/6, 6/7 | | | |
| Visual Count | | | 6/6, 6/7 | | 7/10, 7/11 | 8/4, 8/9 |
| Sticky Card | | 5/29, 5/31 | | 6/30, 6/28 | | 8/1, 8/3 |
| Pitfall Trap | 4/26, 5/1 | 5/29, 5/31 | | 6/30, 6/28 | | 8/1, 8/3 |
| Litter Extraction | 4/26, 5/1 | 5/29, 5/31 | | 6/30, 6/28 | | 8/1, 8/3 |

Table S7. Neonicotinoid residues in soil samples collected before and after soybean was planted in 2015 and maize was planted in 2017. The detection level was 5ppb for imidacloprid (IMI), 10ppb for thiamethoxam (THI) and 15ppb for clothianidin (CLO). Nd = not detected. The pre-planting data for Queenstown for 2015 and 2017 is not presented as no residues were detected in the soil.

| **Crop** | **Site** | **Treatment** | **Insecticide Residue (ppb)** | | | | | |
|---|---|---|---|---|---|---|---|---|
| | | | **Replicate 1 + 2** | | | **Replicate 3 + 4** | | |
| | | | **IMI** | **THI** | **CLO** | **IMI** | **THI** | **CLO** |
| 2015 Soybean Pre-Plant | Beltsville | Control | 8 | nd | Trace | Trace | nd | nd |
| | | Fungicide | 6 | nd | nd | 7 | nd | nd |
| | | Imidacloprid | Trace | nd | nd | Trace | nd | nd |
| | | Thiamethoxam | 7 | nd | Trace | 6 | nd | nd |
| 2015 Soybean Post-Plant | Beltsville | Control | 10 | nd | Trace | Trace | nd | nd |
| | | Fungicide | Trace | nd | nd | 8 | nd | nd |
| | | Imidacloprid | 8 | nd | nd | Trace | Trace | nd |
| | | Thiamethoxam | Trace | nd | Trace | 8 | Trace | nd |
| | Queenstown | Control | nd | nd | nd | nd | nd | nd |
| | | Fungicide | nd | nd | nd | nd | nd | nd |
| | | Imidacloprid | Trace | nd | nd | nd | nd | nd |
| | | Thiamethoxam | nd | 16 | nd | nd | nd | nd |
| 2017 Maize Pre-Plant | Beltsville | Control | 7 | nd | nd | nd | nd | nd |
| | | Fungicide | nd | nd | nd | nd | nd | nd |
| | | Imidacloprid | 8 | nd | nd | 9 | nd | nd |
| | | Thiamethoxam | Trace | nd | nd | nd | nd | nd |
| 2017 Maize Post-Plant | Beltsville | Control | 7 | nd | nd | nd | nd | nd |
| | | Fungicide | Trace | nd | nd | nd | nd | nd |
| | | Imidacloprid | 11 | nd | nd | 35 | nd | nd |
| | | Thiamethoxam | 12 | 17 | 23 | Trace | nd | nd |
| | Queenstown | Control | nd | nd | nd | nd | nd | nd |
| | | Fungicide | nd | nd | nd | nd | nd | nd |
| | | Imidacloprid | 14 | nd | nd | 26 | nd | nd |
| | | Thiamethoxam | nd | 15 | nd | nd | 16 | nd |
| | | | **Replicate 1+2+3+4** | | | | | |
| | | | IMI | THI | CLO | | | |
| 2015-2016 Winter Wheat | Beltsville | Control | Trace | nd | nd | | | |
| | | Fungicide | Trace | nd | nd | | | |
| | | Imidacloprid | 7 | nd | nd | | | |
| | | Thiamethoxam | Trace | nd | nd | | | |
| | Queenstown | Control | nd | nd | nd | | | |
| | | Fungicide | nd | nd | nd | | | |
| | | Imidacloprid | Trace | nd | nd | | | |
| | | Thiamethoxam | nd | nd | nd | | | |

Table S8. Taxa collected through pitfall traps that comprised at least 1% of total abundance in one or more crops Data from subsamples was averaged and data was totaled across locations and sampling dates. Total organisms collected includes ants (Formicidae) and insects from the orders Coleoptera, Diptera, Hemiptera, Hymenoptera and Lepidoptera that could not be identified beyond order, which were excluded from analysis. FS = full season, DC = double cropped.

| **Guild** | **Taxa** | | **% Total** | | | |
|---|---|---|---|---|---|---|
| | | | **2015 FS Soybean** | **2016 DC Soybean** | **2015-2016 Wheat** | **2017 Maize** |
| Predator | Chilopoda | | 0.8 | 0.4 | 1.0 | 0.6 |
| | Araneae | | 9.4 | 2.7 | 7.9 | 3.9 |
| | Mesostigmata | | 2.5 | 1.2 | 5.7 | 1.2 |
| | Coleoptera | Staphylinidae | 4.3 | 3.9 | 3.2 | 2.2 |
| | | Carabidae | 1.6 | 0.6 | 1.3 | 1.5 |
| | | Cantharidae | 5.2 | 0.1 | <0.1 | <0.1 |
| Parasitoid | Hymenoptera | Scelionidae | 0.8 | 1.0 | 0.5 | 0.8 |
| Pest | Gastropoda | | 0.9 | <0.1 | 0.8 | 3.6 |
| | Hemiptera | Pentatomidae *herbivorous* | 2.1 | 0.1 | 0.1 | 2.2 |
| | Odonata | Gryllidae | 2.5 | 4.5 | 0.9 | 5.7 |
| Other | Lumbricina | | 0.3 | <0.1 | 0.7 | 1.2 |
| | Diplopoda | | 0.9 | 0.1 | 1.2 | 0.5 |
| | Acari | Tarsonemidae & Oribatida | 16.9 | 5.1 | 4.4 | 11.1 |
| | Collembola | | 32.1 | 68.4 | 53.8 | 49.0 |
| | Coleoptera | Mycetophagidae | 0.3 | 0.3 | 1.9 | 0.2 |
| | | Cryptophagidae | <0.1 | 0.1 | 2.1 | 0.2 |
| | Diptera | Chloripidae | 3.8 | 0.1 | 0.1 | 0.4 |
| | | Sciaridae | 0.6 | 0.5 | 2.0 | 0.7 |
| | | Phoridae | 0.7 | 1.1 | 1.5 | 0.2 |
| | Hymenoptera | Formicidae | 10.2 | 5.7 | 3.9 | 11.6 |
| **Organisms Analyzed** | | | **9757** | **24768** | **9457** | **9377** |
| **Total Organisms Collected** | | | **11057** | **26549** | **10028** | **10664** |

Table S9. Taxa collected through litter extraction that comprised at least 1% of total abundance in one or more crops. Data from subsamples was averaged and data was totaled across locations and sampling dates. Total organisms collected includes ants (Formicidae) and insects from the orders Coleoptera, Diptera, Hemiptera, Hymenoptera and Lepidoptera that could not be identified beyond order, which were excluded from analysis. FS = full season, DC = double cropped.

| Guild | Taxa | | % Total | | | |
|---|---|---|---|---|---|---|
| | | | 2015 FS Soybean | 2016 DC Soybean | 2015-2016 Wheat | 2017 Maize |
| Predator | Araneae | | 1.2 | 1.6 | 1.5 | 2.1 |
| | Mesostigmata | | 19.6 | 15.0 | 18.8 | 8.1 |
| | Coleoptera | Staphylinidae | 1.5 | 0.5 | 0.9 | 1.6 |
| Pest | Thysanoptera | Thripidae | 0.7 | 2.5 | 2.4 | 0.8 |
| Other | Lumbricina | | 0.7 | 0.4 | 0.5 | 2.3 |
| | Diplopoda | | 0.5 | 0.1 | 0.7 | 1.5 |
| | Acari | Tarsonemidae & Oribatida | 47.5 | 43.8 | 46.4 | 48.9 |
| | Collembola | | 20.5 | 29.2 | 20.6 | 18.2 |
| | Hymenoptera | Formicidae | 1.9 | 1.4 | 0.8 | 5.7 |
| **Organisms Analyzed** | | | **22117** | **23139** | **18536** | **5540** |
| **Total Organisms Collected** | | | **23240** | **24250** | **19499** | **6101** |

Table S10. Taxa collected through sticky cards that comprised at least 1% of total abundance in one or more crops Data from subsamples was averaged and data was totaled across locations and sampling dates. Total organisms collected includes ants (Formicidae) and insects from the orders Coleoptera, Diptera, Hemiptera, Hymenoptera and Lepidoptera that could not be identified beyond order, which were excluded from analysis. Sticky cards from the first double-cropped soybean sampling date at Queenstown were misplaced, and so only data from the second and third dates is included. FS = full-season, DC = double-cropped.

| **Guild** | **Taxon** | | **% Total** | | | |
|---|---|---|---|---|---|---|
| | | | **2015 FS Soybean** | **2016 DC Soybean** | **2015-2016 Wheat** | **2017 Maize** |
| Predator | Coleoptera | Coccinelidae | 0.1 | 0.3 | 0.5 | 1.3 |
| | Hemiptera | Anthocoridae | 1.2 | 0.9 | <0.1 | 4.5 |
| Parasitoid | Hymenoptera | Scelionidae | 1.7 | 3.0 | 4.2 | 2.6 |
| | | Ceraphronidae | 3.8 | 2.4 | 8.5 | 5.0 |
| | | Aphelinidae | 3.4 | 1.6 | 7.6 | 1.5 |
| | | Mymaridae | 1.4 | 3.5 | 2.2 | 4.4 |
| | | Eulophidae | 0.1 | 0.2 | 3.2 | 0.6 |
| | | Trichogrammatidae | 0.4 | 1.5 | 0.6 | 1.1 |
| | | Braconidae | 0.3 | 1.0 | 3.5 | 0.6 |
| | Diptera | Tachinidae | 0.7 | 1.1 | 0.6 | 0.9 |
| Pest | Coleoptera | Chrysomelidae - Alticini | 0.4 | 0.8 | 0.5 | 2.4 |
| | Hemiptera | Cicadellidae | 7.5 | 16.5 | 8.3 | 16.0 |
| | | Aphididae | 1.8 | 1.8 | 1.8 | 4.2 |
| | | Aleyrodidae | <0.1 | 2.0 | 0.4 | 0.7 |
| | Hymenoptera | Cynipidae | 0.3 | 2.4 | 1.4 | 1.6 |
| | Thysanoptera | Thripidae | 32.7 | 10.2 | 31.8 | 12.5 |
| Other | Coleoptera | Phalacridae | 0.1 | 0.1 | 0.9 | 2.7 |
| | Diptera | Chloropidae | 32.4 | 38.3 | 10.3 | 20.2 |
| | | Sciaridae | 2.0 | 2.5 | 3.1 | 4.5 |
| | | Phoridae | 1.0 | 0.9 | 0.5 | 0.1 |
| | | Cecidomyiidae | 0.8 | 2.8 | 3.5 | 7.1 |
| **Organisms Analyzed** | | | **13997** | **9799** | **5284** | **5333** |
| **Total Organisms Collected** | | | **14034** | **9906** | **5303** | **5354** |

Table S11. Taxa collected through sweep net sampling that comprised at least 1% of total abundance in soybean in 2015 or 2016. Data was totaled across locations. The percent total is included for each group, as well as the overall abundance for that crop. Total organisms collected includes insects from the orders Coleoptera, Diptera, Hemiptera, Hymenoptera and Lepidoptera that could not be identified beyond order, which were excluded from analysis. Samples from one imidacloprid replicate at the Beltsville, MD site were misplaced prior to processing in 2015.

| **Guild** | **Taxa** | | **% Total** | |
|---|---|---|---|---|
| | | | **2015 Soybean** | **2016 DC Soybean** |
| Predator | Araneae | | 8.6 | 7.2 |
| | Coleoptera | Coccinelidae | 0.6 | 1.7 |
| | Hemiptera | Anthocoridae | 11.7 | 5.2 |
| | | Geocoridae | 0.3 | 1.3 |
| | | Nabidae | 0.3 | 1.0 |
| Parasitoid | Hymenoptera | Braconidae | 0.6 | 2.4 |
| | | Scelionidae | 1.3 | 0.8 |
| Pest | Coleoptera | Misc. Chrysomelidae | 1.0 | 0.0 |
| | | Chryosmelidae - Alticini | 0.2 | 1.1 |
| | | Curculionidae | 3.7 | 0.0 |
| | | *Diabrotica/Acalymma* spp. | 1.7 | 1.8 |
| | | *Epilachna varivestis* | 1.3 | 0.0 |
| | | Scarabaeidae | 1.7 | 2.9 |
| | Hemiptera | Aphididae | 2.9 | 0.5 |
| | | Cicadellidae | 3.6 | 5.9 |
| | | Pentatomidae herbivorous | 2.5 | 0.3 |
| | Lepidoptera | *Hypena scabra* | 3.1 | 18.2 |
| | | *Spodoptera frugiperda* | 0.0 | 4.6 |
| | Orthoptera | Acrididae | 0.0 | 2.4 |
| Other | Diptera | Chloropidae | 46.6 | 31.3 |
| | | Phoridae | 0.4 | 1.0 |
| | | Sciaridae | 0.0 | 1.6 |
| | Hymenoptera | Formicidae | 0.2 | 1.3 |
| **Organisms Analyzed** | | | **2320** | **1572** |
| **Total Organisms Collected** | | | **2358** | **1558** |

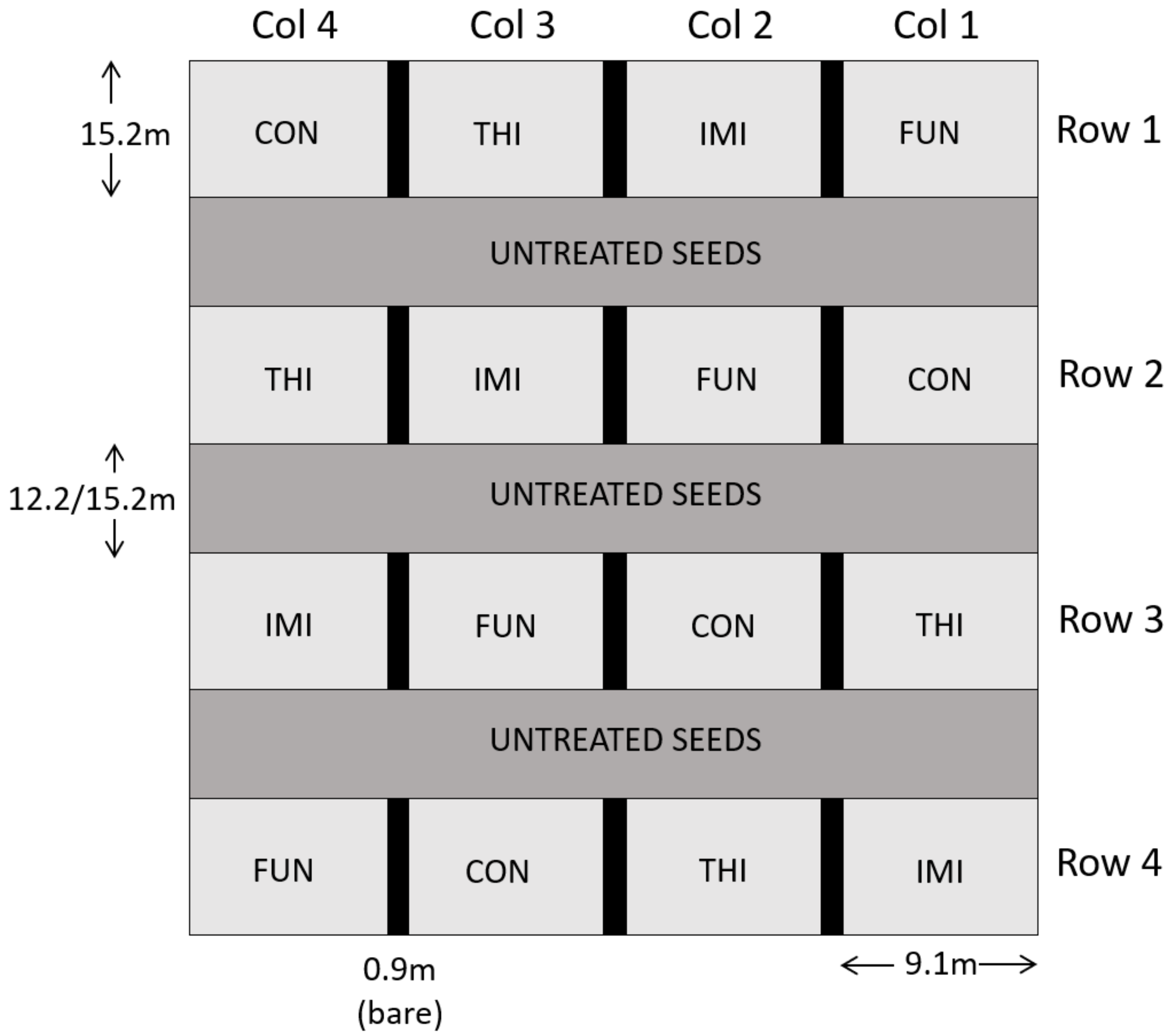


Fig. S1. Plot map showing the Latin square arrangement of four replicates of each treatment [control (CON), fungicide only (FUN), imidacloprid + fungicide (IMI), thiamethoxam + fungicide (THI)]. Rows were separated by turn rows planted with untreated grain (12.2m at Queenstown, 15.2m at Beltsville), and columns were separated by bare strips.

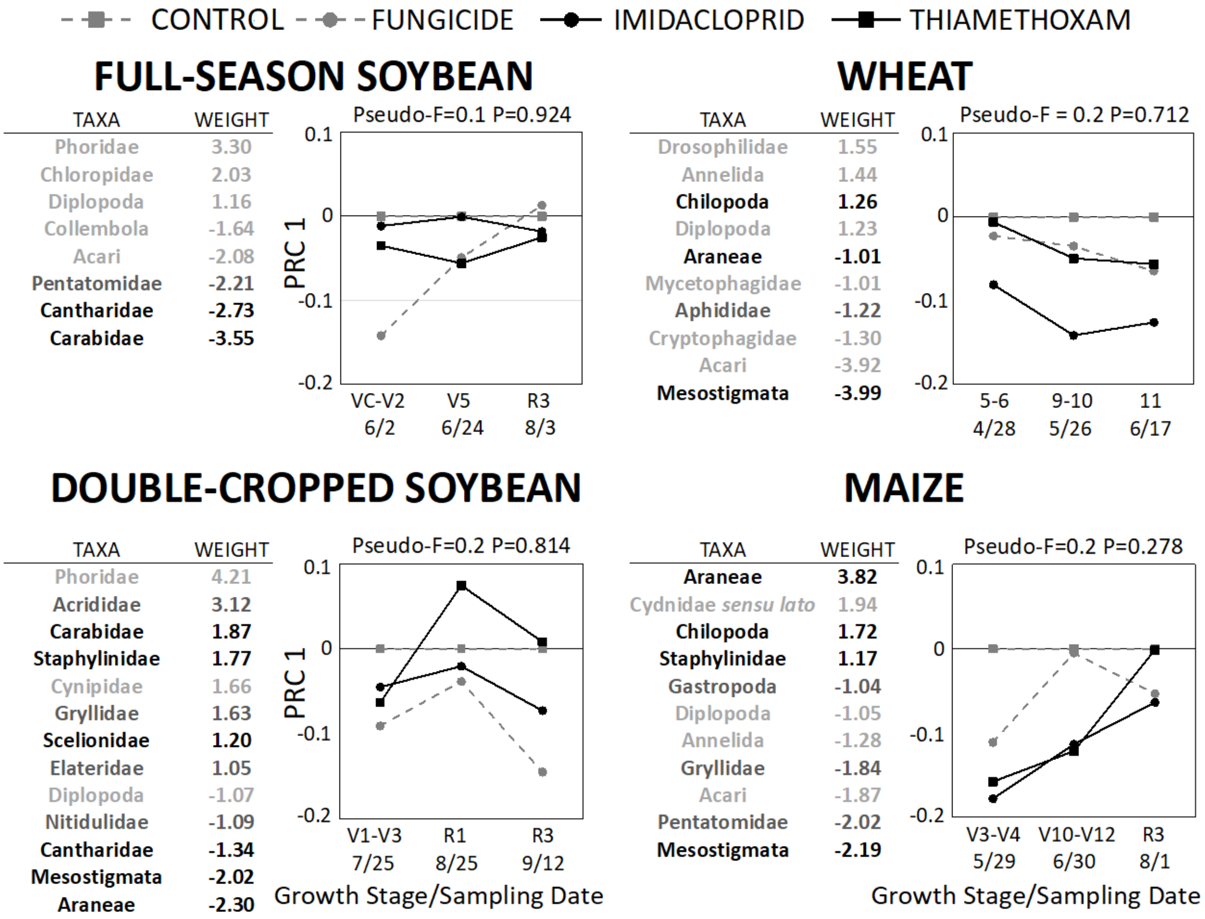


Fig. S2. Principal Response Curve analysis of pitfall trap data for all crops. For each crop, date*treatment served as the explanatory variable, with date and site*replicate used as covariates. Subsamples were averaged for each replicate, and only taxa with overall means greater than one were included. Ants (Formicidae) were also excluded due to their highly clumped distribution. A Monte-Carlo permutation procedure with N=499 was used to calculate the Pseudo-F statistic. Taxon weights indicate which groups most contributed to the observed community response. Higher positive weights indicate that taxon abundances in the treated plots followed the trend depicted by the response curve, whereas higher negative values indicate the opposite. Taxon weights between -1 and 1 were excluded due to weak response or lack of correlation with the trends shown. Beneficial groups are shown in black, economic pests in dark gray, and other groups in light gray. Acari refers specifically to the mite order Oribatida and the family Tarsonemidae.

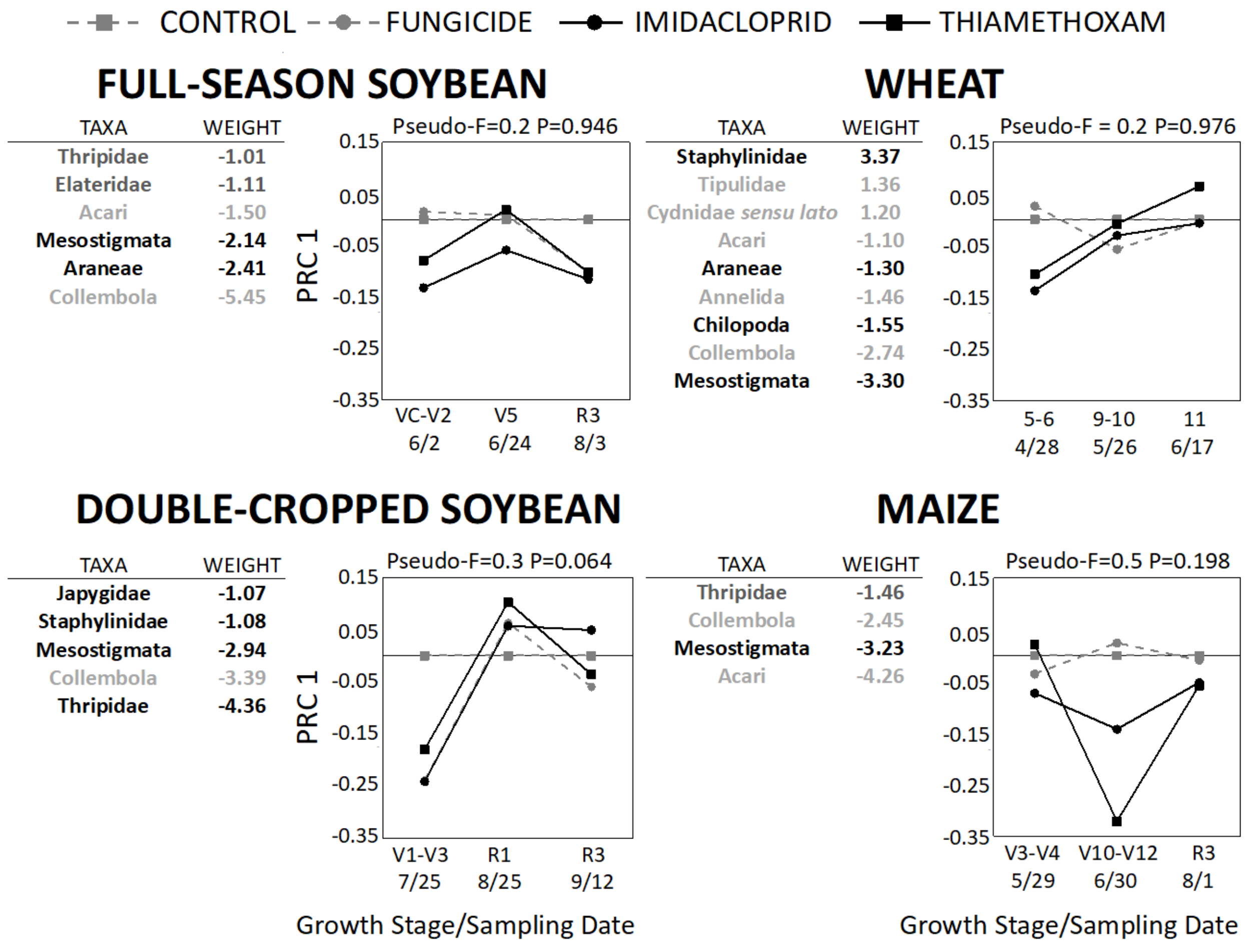


Fig. S3. Principal Response Curve analysis of litter extraction data for all crops. For each crop, date*treatment served as the explanatory variable, with date and site*replicate used as covariates. Subsamples were averaged for each replicate, and only taxa with overall means greater than one were included. Ants (Formicidae) were also excluded due to their highly clumped distribution. A Monte-Carlo permutation procedure with N=499 was used to calculate the Pseudo-F statistic. Taxon weights indicate which groups most contributed to the observed community response. Higher positive weights indicate that taxon abundances in the treated plots followed the trend depicted by the response curve, whereas higher negative values indicate the opposite. Taxon weights between -1 and 1 were excluded due to weak response or lack of correlation with the trends shown. Beneficial groups are shown in black, economic pests in dark gray, and other groups in light gray. Acari refers specifically to the mite order Oribatida and the family Tarsonemidae.
